# Expanding the biotechnological scope of metabolic sensors through computation-aided designs

**DOI:** 10.1101/2024.08.23.609350

**Authors:** Enrico Orsi, Helena Schulz-Mirbach, Charles A. R. Cotton, Ari Satanowski, Henrik M. Petri, Susanne L. Arnold, Natalia Grabarczyk, Rutger Verbakel, Karsten S. Jensen, Stefano Donati, Nicole Paczia, Timo Glatter, Andreas M. Küffner, Tanguy Chotel, Farah Schillmüller, Alberto De Maria, Hai He, Steffen N. Lindner, Elad Noor, Arren Bar-Even, Tobias J. Erb, Pablo I. Nikel

## Abstract

Metabolic sensors are microbial strains modified so that biomass formation correlates with the availability of specific metabolites. These sensors are essential for bioengineering (e.g. in growth-coupled designs) but creating them is often a time-consuming and low-throughput process that can potentially be streamlined by *in silico* analysis. Here, we present the systematic workflow of designing, implementing, and testing versatile *Escherichia coli* metabolic sensor strains. Glyoxylate, a key metabolite in (synthetic) CO_2_ fixation and carbon-conserving pathways, served as the test molecule. Through iterative screening of a compact metabolic model, we identified non-trivial growth-coupled designs that resulted in six metabolic sensors with a wide sensitivity range for glyoxylate, spanning three orders of magnitude in detected concentrations. We further adapted these *E. coli* strains for sensing glycolate and demonstrated their utility in both pathway engineering (testing a key metabolic module *via* glyoxylate) and applications in environmental monitoring (quantifying glycolate produced by photosynthetic microalgae). The versatility and ease of implementation of this workflow make it suitable for designing and building multiple metabolic sensors for diverse biotechnological applications.

**Teaser:** A streamlined workflow enables the rapid design of versatile *E. coli* metabolic sensors for detecting key metabolites in bioengineering and monitoring.

## Introduction

Metabolism is the set of chemical reactions sustaining life (Downs, 2006). These reactions can be systematically investigated thanks to standardized approaches in synthetic biology, a field which applies engineering principles to living (micro)organisms (Decoene et al., 2018; Lv et al., 2022). Microbial auxotrophic strains with defined disruptions of the metabolic network can be designed to have a defined demand for a specific metabolite and thus are ideal platforms for metabolic studies (Orsi et al., 2021; Wenk et al., 2018). Given their dependence on given metabolic intermediates, we will refer to these selection strains as metabolic sensors. Originally used for biochemistry studies through growth complementation, metabolic sensors are now increasingly used for studying metabolic pathways *in vivo* (Chen et al., 2020; Gleizer et al., 2019; Kim et al., 2020; Schulz-Mirbach et al., 2022; Wenk et al., 2022).

Construction of metabolic sensor strains requires reliable quantitative predictions of the effect of gene deletions which must be confirmed *in vivo*. Yet the implementation of auxotrophic phenotypes still relies on one-at-a-time interventions (e.g. multiple rounds of deletion and testing) with the exception of a handful of cases where systematic analysis of all the possible options was performed through screening of metabolic models (Niv Antonovsky et al., 2016; Kim et al., 2020; Satanowski et al., 2020). In all these cases, computations were performed on a core metabolic model (limited to glycolysis/gluconeogenesis, pentose phosphate pathway, tricarboxylic acid cycle, and oxidative phosphorylation), where the output is inherently limited to growth-coupled designs around the 12 universal biomass precursors belonging to the central carbon metabolism (Neidhardt et al., 1990). However, several additional reactions are involved in the synthesis of key biomass precursors *in vivo* (e.g. amino acids or lipids biosynthesis), hence larger metabolic models could lead to multiple growth-coupled designs that can be further explored. However, a model encompassing the entirety of an organism’s metabolism is highly complex, which could increase computing times and lead to irrelevant or unfeasible solutions based on secondary reactions. A compromise between these two approaches would be to employ a medium-scale metabolic model covering core metabolism and essential metabolic pathways (i.e. the synthesis energy carriers and biosynthetic precursors like amino acids) without the redundancies of a genome-scale model (Corrao et al., 2024).

In the work presented here we designed metabolic sensors for glyoxylate, a non-essential metabolite which is not directly involved in the synthesis of biomass precursors in *Escherichia coli*. Creating metabolic sensors for this molecule therefore requires deep metabolic rewiring. We adopted a medium-scale metabolic model to guide the design and engineering of several metabolic sensors for this metabolic hub which was studied in the context of prebiotic chemistry (Krishnamurthy & Liotta, 2023; Pulletikurti et al., 2022; Springsteen et al., 2018; Yadav et al., 2022), synthetic metabolism (Mainguet et al., 2013; McLean et al., 2023; Ren et al., 2018; Schwander et al., 2016), and biomanufacturing (Vuoristo et al., 2015; Yang et al., 2022). We show that complex screening applications can be accessed using our sensor strains. These include *in vivo* screening of enzyme variants for synthetic one-carbon (C1) assimilation and sensing of excreted glycolate produced during photorespiration as a globally occurring environmental process. This work improves the robustness of the workflow for computational design of metabolic interventions and yields an array of novel glyoxylate and glycolate sensors supporting applications beyond the traditional uses of metabolic sensors.

## Results

### Systematic interrogation of a compact metabolic reconstruction for growth-coupled designs around glyoxylate

Since glyoxylate is not an essential metabolite for wild-type *E. coli,* coupling this organism’s growth to glyoxylate availability requires complex metabolic rewiring. Therefore, we propose a pipeline which predicts potential designs *in silico* using a compact metabolic model (**Fig. 1A**). To systematically investigate all the combinations of knockouts (KOs) that force growth to be dependent on glyoxylate, we used an algorithm we had previously applied on a core *E. coli* model (Niv Antonovsky et al., 2016; Aslan et al., 2020), but replaced the core-model with the recently published iCH360 medium-scale model of *E. coli’*s metabolism (Corrao et al., 2024). This medium-scale model (323 reactions) offers wider coverage of possible KOs compared to a core model (95 reactions) and limits the number of additional experience-based manual interventions required compared to a full genome-scale model (2000 reactions or above). To adapt the medium-scale model to identify growth-coupled designs for glyoxylate, we included four additional reactions: (i) glyoxylate uptake; (ii) an aspartate-glyoxylate aminotransferase (BHC) (Schada Von Borzyskowski et al., 2019) as proxy for promiscuous transaminase activities on glyoxylate; (iii) glyoxalate carboligase (GLXCL); and (iv) tartronate semialdehyde reductase (TRSARr) (**Fig. 1B**). The procedure then iterates through a list of KO combinations and uses flux balance analysis to test which combinations cause growth to be dependent on glyoxylate using different carbon substrates (Niv Antonovsky et al., 2016; Aslan et al., 2020).

**Fig. 1.**
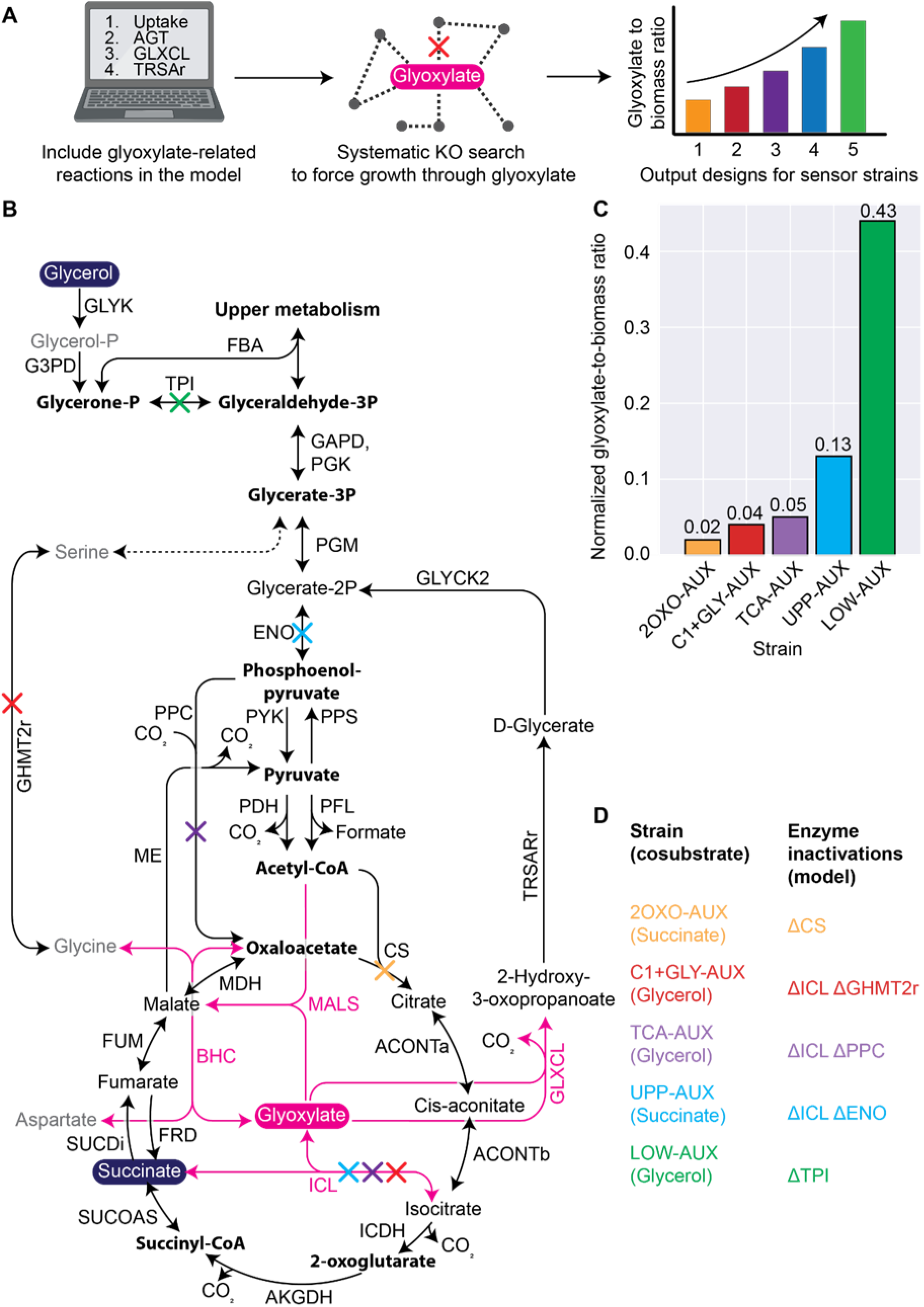
*In silico* modeling for exploring glyoxylate-dependent growth-coupling schemes. (**A**) The workflow we followed started by adapting a medium-scale metabolic model of *Escherichia coli* by including reactions involved in glyoxylate production or consumption: glyoxylate uptake, BHC (aspartate-glyoxylate aminotransferase), GLXCL (glyoxalate carboligase), TRSAr (tartronate semialdehyde reductase). Next, the updated model was used to run the algorithm for systematically determining combinations of gene deletions that require glyoxylate for growth. As output, the algorithm proposes different combinations of such inactivations and ranks them quantitatively in terms of glyoxylate demand (represented by a “glyoxylate-to-biomass ratio”, GBR). (**B**) Schematic overview of the central carbon metabolism of *E. coli* covered by the adapted model. Glyoxylate is shown in magenta together with the reactions (and associated enzymes) originating from this molecule. Succinate and glycerol are shown (navy blue) because these metabolites are chosen as added carbon sources for the design of individual glyoxylate sensor strains. Essential biomass precursors are indicated in bold. Enzymes are abbreviated following BiGG nomenclature. Colored arrows indicate reactions deleted in certain sensor strain designs as proposed by the algorithm to force glyoxylate-dependence. Each color represents a different sensor strain design. (**C**) Graphic output highlighting the different combinations of enzyme inactivations proposed by the algorithm. The normalized GBR value is a unitless value obtained by dividing the GBR of the sensor by the GBR of the wild-type strain growing on glyoxylate as unique carbon source. (**D**) Legend of strains and associated enzyme inactivations selected from the algorithm.

With this extended medium-scale model, we iteratively explored how different combinations of gene deletions affect the dependency of *E. coli* growth on glyoxylate. We set a maximum of two KOs to force glyoxylate consumption for biomass buildup. To broaden the spectrum of identified designs, we performed this search for growth on two different main carbon sources (glycerol and succinate) not affected by catabolite repression (**Fig. 1B**). As output, the algorithm provided a set of different KO combinations for each of the two carbon sources, spanning different levels of dependency on glyoxylate (**Supplementary Figure 1**).

From these results, we decided to pursue five designs covering different glyoxylate-derived biomass precursors and glyoxylate-demands, namely the amount required for the synthesis of biomass (**Fig. 1C**). The first group included disruptions of glycolysis/gluconeogenesis, causing reliance on glyoxylate either for the synthesis of intermediates in upper glycolysis (e.g. sugar phosphates, “upper metabolism”) or for the synthesis of “lower metabolism”. The second group included deletions requiring glyoxylate for anaplerosis of the TCA cycle. The third group included designs which require glyoxylate for glycine biosynthesis through transamination. Notably, this last group of designs would not have been accessible by use of a core model of *E. coli* metabolism because key metabolic routes of amino acid metabolism were not included i.e., serine and glycine interconversion.

The algorithm provided the minimum set of KOs required for generating each auxotrophic phenotype in the metabolic sensors. However, since this *in silico* prediction does not consider factors such as metabolic regulation and enzyme promiscuities, it needs to be complemented by literature knowledge of the host’s metabolic network and physiology. Thus, we manually expanded the knockout selection with additional targets known from literature to support the strain engineering step (**Fig. 2A**). In the next section, we describe how we moved from the *in silico* predictions to successfully engineering the respective metabolic sensors.

**Fig. 2.**
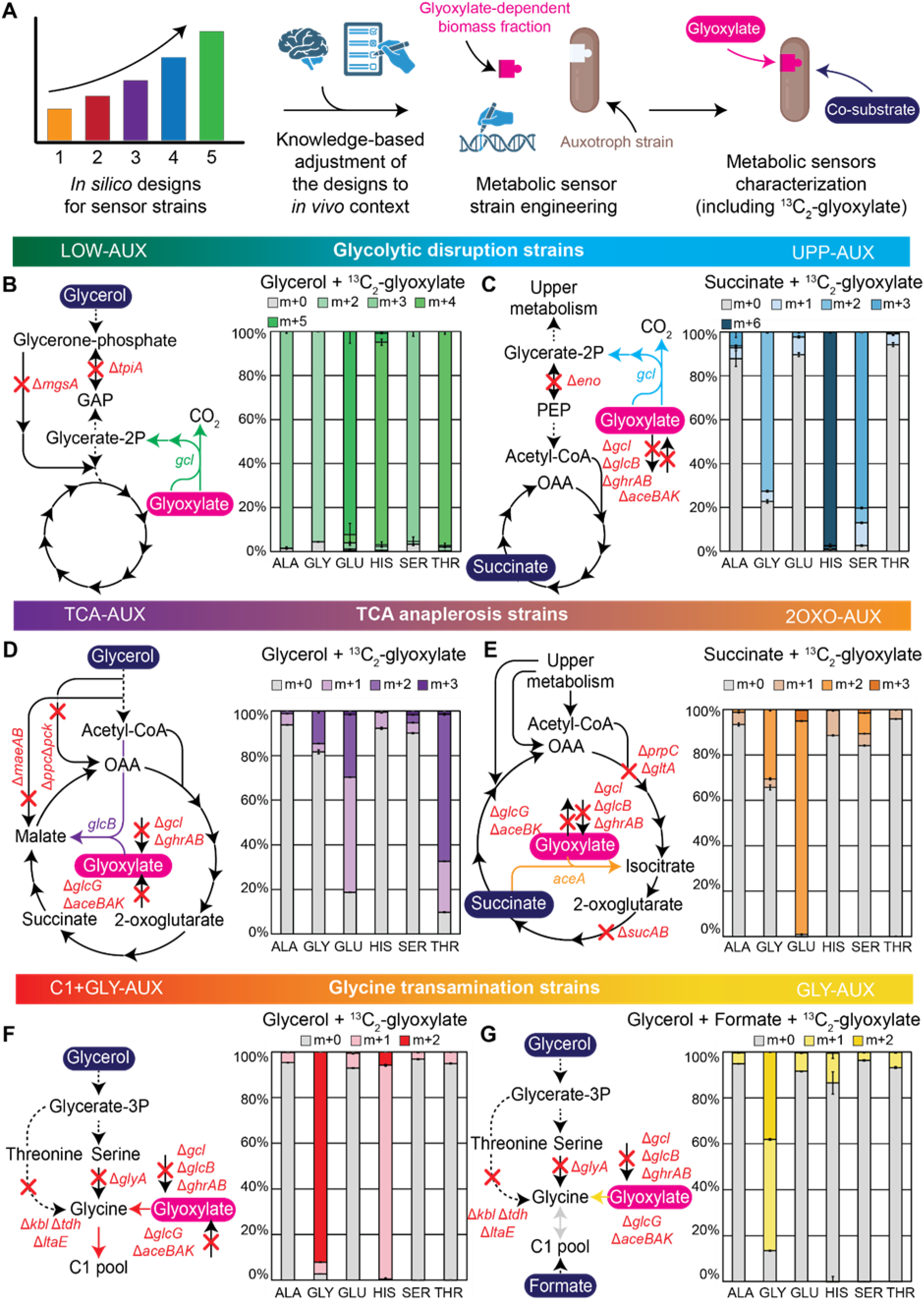
Metabolic sensor strains engineering and characterization. (**A**) The workflow for creating the metabolic sensors starts with the output from the algorithm, which is complemented by curated literature knowledge on the host’s metabolic network. The strains are engineered accordingly, and the auxotrophic phenotype for the target molecule (glyoxylate) is characterized and confirmed. (**B-G**) overview of the metabolic sensors and corresponding ^13^C-labeled amino acids pattern (one letter code used) as a result of cultivation with ^13^C_2_-glyoxylate. Abbreviations for the genes *mgsA*: methylglyoxal synthase; *tpiA*: triose-phosphate isomerase; *eno*: enolase; *gcl*: glyoxylate carboligase; *glcB*: malate synthase G; *ghrA*: glyoxylate/hydroxypyruvate reductase A; *ghrB*: glyoxylate reductase; *aceA*: isocitrate lyase; *aceB*: malate synthase A; *aceK*: isocitrate dehydrogenase kinase / isocitrate dehydrogenase phosphatase; *maeA*: malate dehydrogenase (oxaloacetate-decarboxylating); *maeB*: malate dehydrogenase (oxaloacetate-decarboxylating) (NADP^+^); *ppc*:phosphoenolpyruvate carboxylase; *pck*: phosphoenolpyruvate carboxykinase (ATP); *gltA*: citrate synthase; *prpC*: 2-methylcitrate synthase; *sucA*: 2-oxoglutarate decarboxylase, thiamine-requiring; *sucB*: 2-oxoglutarate dehydrogenase E2 subunit; *kbl*: 2-amino-3-ketobutyrate CoA ligase; *tdh*: threonine dehydrogenase; *ltaE*: low-specificity L-threonine aldolase; *glyA*: serine hydroxymethyltransferase.

### Metabolic sensor engineering and corresponding glyoxylate contribution to biomass buildup

We started by engineering the group of metabolic sensors that rely on disruptions of glycolysis/gluconeogenesis. The selection predicted to have the highest glyoxylate demand is based on inactivation of triose phosphate isomerase (encoded by the gene *tpiA*) (**Fig. 2B**). Yet this deletion alone is known to be bypassed during growth on glycerol by activation of the methylglyoxal pathway (Iacometti et al., 2022; Long & Antoniewicz, 2019b). To avoid this bypass, we additionally deleted *mgsA* (encoding methylglyoxal synthase), thus generating a double mutant Δ*tpiA* Δ*mgsA* (Iacometti et al., 2022). We refer to this strain as LOW-AUX because it needs glyoxylate to supplement all metabolism downstream of glyceraldehyde 3-phosphate (“lower metabolism”). After validating the sensor’s dependency on glyoxylate, we confirmed the expected metabolic fluxes by tracing ^13^C incorporation into proteinogenic amino acids during growth of the LOW-AUX strain on uniformly labelled ^13^C_2_-glyoxylate, with unlabeled glycerol serving as the main carbon source. The resulting data confirmed labeling incorporation in the majority of the proteinogenic amino acids, thereby demonstrating contribution of glyoxylate to the buildup of a large fraction of biomass precursors (**Fig. 2B, Supplementary Figure 2, Supplementary Figure 3A**). The other glycolytic disruption requires the combined inactivation of enolase (ENO, encoded by *eno*) and isocitrate lyase (ICL, encoded by *aceA*). This strain relies on succinate supplementation to feed “lower metabolism”, while supplied glyoxylate should be converted to glycerate 2-phosphate, feeding “upper metabolism” (therefore named UPP-AUX). To prevent futile consumption of glyoxylate via the glyoxylate shunt, we decided to delete the relevant reactions. These additional deletions included *aceB* and *glcB* (malate synthases) as well as *ghrA* and *ghrB* (glyoxylate/hydroxypyruvate reductases) (**Fig. 2C**). In this strain, glyoxylate followed a different fate which was traced using ^13^C labeling, confirming its contribution to the synthesis of amino acids derived from upper metabolism (**Fig. 2C, Supplementary Figure 2, Supplementary Figure 3B**).

Then, we implemented the designs interrupting TCA cycle anaplerosis. We started with a strain including inactivations of PEP carboxylase (*ppc*), PEP carboxykinase (*pckA*) and isocitrate lyase (*aceA*) (Claassens et al., 2020). In addition, the strain included adaptations to streamline glyoxylate utilization through constitutive, chromosomal overexpression of malate synthase (*glcB*, **Fig. 2D**). Moreover, we deleted *maeA* and *maeB* (encoding two malic enzymes, both canonically involved in gluconeogenesis but thermodynamically reversible under certain conditions), Δ*aceB* (additional copy of malate synthase) and Δ*aceK* (regulator of isocitrate dehydrogenase), Δ*gcl* (glyoxalate carboligase, deleted to prevent wasteful conversion of glyoxylate through tartronate semialdehyde), and Δ*ghrA* and *ΔghrB* (glyoxylate/hydroxypyruvate reductase). Therefore, this corresponding TCA-AUX strain relies on malate synthase activity for the condensation of glyoxylate and acetyl-CoA for the biosynthesis of all TCA cycle intermediates (**Fig. 2D**). Labelling patterns in the TCA-AUX grown with ^13^C_2_-glyoxylate and unlabeled glycerol reflect the expected fluxes, with amino acids derived from TCA cycle intermediates predominantly labelled (**Fig. 2D, Supplementary Figure 2, Supplementary Fig. 3C**).

The other TCA cycle-related design, which involved inactivation of citrate synthase, was realized by deleting *gltA* (citrate synthase) and *prpC* [2-methylcitrate synthase, reported to have promiscuous citrate synthase activity (Gerike et al., 1998; Guzmán et al., 2015)] (**Fig. 2E**). In addition to these deletions, we aimed to reduce the demand for glyoxylate by deleting *sucAB* (2-oxoglutarate dehydrogenase), thereby limiting the glyoxylate requirement to the synthesis of the biomass precursor 2-oxoglutarate (2OXO-AUX strain). This strain required additional supplementation of succinate to prevent succinyl-CoA auxotrophy (**Supplementary Figure 5**). To prevent undesired glyoxylate consumption through other routes we abolished suspected glyoxylate sinks by adding further deletions as in the TCA-AUX, i.e., Δ*gcl*, Δ*ghrAB*, Δ*glcB*, and Δ*aceBK*. Isotopic ^13^C labeling patterns of the 2OXO-AUX strain growing with ^13^C_2_-glyoxylate and unlabeled succinate confirmed incorporation of glyoxylate into 2-oxoglutarate with 100% of glutamate being labeled twice (m+2), while the other TCA cycle-derived amino acids aspartate, lysine and threonine remained unlabeled (m+0) (**Fig. 2E, Supplementary Figure 2, Supplementary Figure 4A**).

In the third and last selection strategy, we pursued transamination of glyoxylate for synthesis of the essential biomass precursor glycine. Glycine is also involved in the synthesis of the essential C1 metabolites [5,10-methylene-tetrahydrofolate (methylene-THF) and 10-formyl-THF]. Thus, we envisioned two setups for glyoxylate sensor strains depending on inclusion or not of the C1 pool, named C1+GLY-AUX and GLY-AUX strains, respectively (**Fig. 2F, G**). First, we implemented deletions that force glyoxylate conversion to glycine through transamination (Δ*gcl* Δ*ghrAB* Δ*glcB*, and Δ*aceBAK*). Second, we prevented glycine synthesis from glycerol by deleting *glyA* (serine hydroxymethyltransferase), *kbl* (2-amino-3-ketobutyrate CoA ligase), and *ltaE* [low-specificity L-threonine aldolase (Liu et al., 1998)]. We demonstrated that this C1+GLY-AUX strain grew in a glyoxylate-dependent manner, thus suggesting that one of the endogenous transaminases is employed for glyoxylate conversion to glycine. Indeed, isotopic labelling patterns of C1+GLY-AUX grown with ^13^C_2_-glyoxylate and unlabeled glycerol confirmed the ability to support glycine synthesis from glyoxylate by showing 100% of the glycine pool being labeled twice (m+2) (**Fig. 2F, Supplementary Figure 2, Supplementary Figure 4B**). Moreover, we proved the glyoxylate contribution to the synthesis of the C1 pool by measuring 100% of the histidine pool with one label (C1 moieties are involved in the synthesis of this amino acid (Yishai et al., 2017)). To gain potential insights into native (promiscuous) transaminases involved in the conversion of glyoxylate to glycine, we searched for differences in protein abundances between the C1+GLY-AUX and a wild type strain grown either with 20 mM glycerol + 1 mM glyoxylate or 20 mM glycerol + 5 mM glycine (**Supplementary Figure 6**). Here, we found that of all known > 20 *E. coli* transaminases, *hisC* (encoding histidinol-phosphate transaminase) was the only one that was significantly upregulated in both comparisons of C1+GLY-AUX versus the wild-type strain on 20 mM glycerol + 1 mM glyoxylate and of C1+GLY-AUX grown with glyoxylate versus with glycine. Additionally, we observed an upregulation of allantoin metabolism genes (*allB* and *allC*) in C1+GLY-AUX grown with glyoxylate in all comparisons, which is in line with previous reports of glyoxylate-dependent expression of these genes (Cusa et al., 1999; Rintoul et al., 2002). While the degradation of the purine synthesis intermediate allantoin proceeds via a ureidoglycine:glyoxylate transaminase (PucG from *Bacillus subtilis* or HpxJ from *Klebsiella pneumoniae*) in other species, *E. coli* MG1655 is not known to harbor a homologue. However, when we used the DELTA-BLAST domain search and PucG from *Bacillus subtilis* (Uniprot ID A0A6M4JLB6) as template, the histidinol-phosphate transaminase (HisC), which was upregulated in our proteomics data, was amongst the hits with the highest identity score (18.32% identity, e-value 1e^-17^). While these findings hint towards relevance of HisC for glyoxylate use in the C1+GLY-AUX, it remains to be investigated whether HisC can promiscuously interconvert glyoxylate and glycine using ureidoglycine as amine donor.

To further boost selection sensitivity, we expressed in the GLY-AUX strain a dedicated aspartate:glyoxylate transaminase (*bhcA*) from *Paracoccus denitrificans* for aiding conversion of glyoxylate to glycine (Schada Von Borzyskowski et al., 2019). Moreover, we interrupted glycine conversion to methylene-THF through deletion of the glycine cleavage system (Δ*gcvTHP*) in the GLY-AUX strain and engineered methylene-THF synthesis from additionally supplied formate by integrating module 1 of the reductive glycine pathway (Yishai et al., 2018) into the chromosome. As we see in the next section, these modifications will impact the strain’s sensitivity to glyoxylate. Remarkably, the strain showed a reduced pool of labeled glycine of almost an equal amount between m+1 and m+2 pools, thereby suggesting a partial contribution of formate to the synthesis of glycine through the reverse glycine cleavage system. Moreover, release of glyoxylate contribution to histidine was confirmed by observing a drastic change in the histidine labeling pattern, with the majority of this amino acid pool remaining unlabeled (m+0) (**Fig. 2G, Supplementary Figure 2, Supplementary Figure 4C**). Once we confirmed that all strains required glyoxylate for the synthesis of one or more essential biomass components, we progressed further into their quantitative characterization.

### Metabolic sensors can detect glyoxylate over a wide concentration range spanning three orders of magnitude

We further analyzed the metabolic sensors by monitoring their growth with varying supplemented concentrations of glyoxylate (**Fig. 3A**). This approach is routinely used in the characterization of metabolic sensors (Aslan et al., 2020; Schulz-Mirbach et al., 2022; Wenk et al., 2020) and allows the range of direct correlation between the concentration of the limiting substrate (glyoxylate) and the final biomass concentration to be identified (**Fig. 3B**). This is determined by plotting the maximal optical density (maxOD_600_) as a function of the concentration of the target metabolite. For our sensitivity analysis, we focused on an operational range giving linear response between substrate concentration and maxOD_600_, i.e., excluding those concentrations where glyoxylate is no longer the limiting metabolite (upper limit) and those were we observed lack of growth because the concentration of glyoxylate is too low (lower limit). We confirmed that the experimentally determined glyoxylate demand was qualitatively consistent with the sensitivity ranking based on predicted “glyoxylate-to-biomass ratios” (GBR) calculated beforehand in flux balance analysis (**Fig. 3B**). In combination, the different constructed metabolite sensor strains enabled a glyoxylate sensitivity range covering three orders of magnitude from 10 µM to 20 mM of glyoxylate (**Fig. 3B**). We confirmed robustness in auxotroph phenotypes for all strains by ruling out any unexpected growth behavior over several days of incubation (**Supplementary Figures 7 and 8**). The upper limit of detection was determined by growing a wild-type strain with glyoxylate serving as the only carbon source, i.e. representing the highest achievable glyoxylate demand (**Supplementary Fig. 9**).

**Figure 3.**
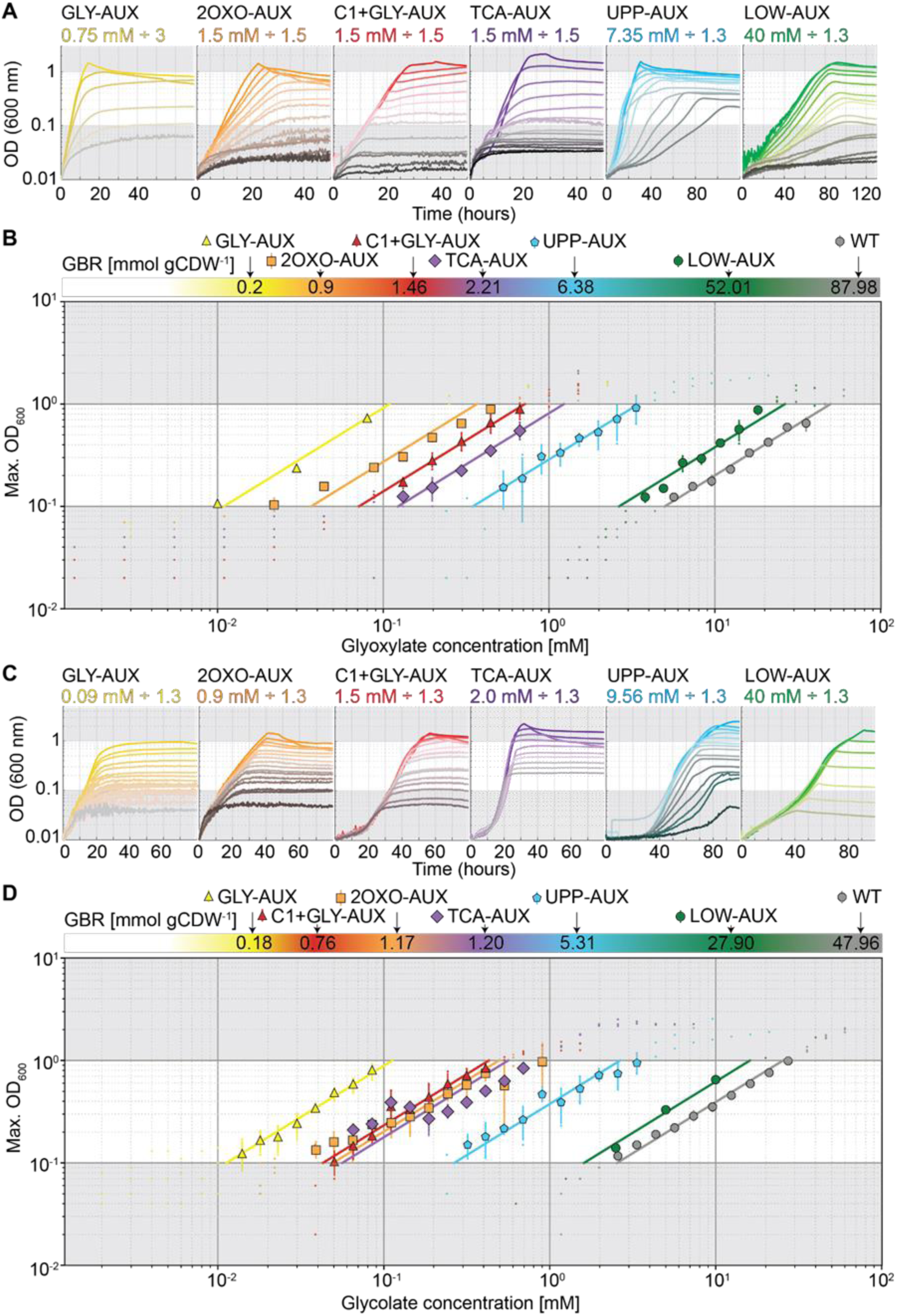
Growth profiles and dose response relationship in the metabolic sensors. (**A**) Growth profiles of the different metabolic sensors with varying concentrations of glyoxylate supplemented in the growth medium. For each strain, the highest concentration of glyoxylate was reported, together with the dilution factor used. (**B**) Relative distribution of the glyoxylate-to-biomass ratio (GBR) for the different strains, calculated by flux balance analysis using the medium-scale model, and correlation between the maximal OD_600_ measured and the initial glyoxylate concentration in the medium. This relationship highlights the linear trend of dose response to glyoxylate at OD_600_ between 0.1 and 1.0. (**C, D**) Same experiments as in A and B but focused on glycolate as target metabolite.

Notably, in some strains the growth rate varied depending on supplied glyoxylate concentrations. This was most likely caused by factors affecting key limiting enzymes required for growth. For example, C1+GLY-AUX and GLY-AUX rely on different transamination reactions to recover growth. Consequently, their growth rate differed significantly (**Fig. 3A, Supplementary Fig. 6**). The faster growth of C1+GLY-AUX was probably due to an increased rate of glycine biosynthesis from glyoxylate caused by the additional engineered expression of a heterologous transaminase (*bhcA*). We could confirm this hypothesis by observing an improved growth rate of the GLY-AUX strain when BhcA was additionally produced in this strain background (**Supplementary Fig. 7**).

### A functional glycolate dehydrogenase complex extends sensors’ detecting ability to glycolate

Once the ability of the sensors to detect glyoxylate was validated, we aimed to extend the sensing ability of the strains to glycolate. The glycolate dehydrogenase complex (GlcDEF) catalyzes the oxidation of glycolate to glyoxylate and is native to *E. coli*. Due to this activity, we expected glycolate to be converted to glyoxylate for subsequent formation of the selected biomass precursors depending on the metabolic context of each sensor strain. Accordingly, when cultivated on glycolate the strains showed comparable growth dependencies as observed with glyoxylate (**Fig. 3C**). Moreover, when plotting the maximum OD_600_ in response to the glycolate concentration provided, the sensor strains could cover a range of concentrations of three orders of magnitude (from 10µM to 20 mM) (**Fig. 3D**). As had been done for glyoxylate, the maximum glycolate detection capacity was determined by growing the wild-type strain on glycolate as carbon source (**Supplementary Fig. 8**).

Altogether, the set of metabolic sensors showed a strict dependence on glyoxylate or glycolate for growth, a dependency that can be exploited to couple growth to modules forming either molecule. In the next sections we showcase two exemplary applications of these metabolic sensors for the screening of enzymatic activities *in vivo* or for the measurement of extracellular glycolate in spent cultivation media.

### *In vivo* testing of malate thiokinase and malyl-CoA lyase through glyoxylate sensing

Next we investigated the use of our metabolic sensors for prototyping metabolic pathways (**Fig. 4A**). A versatile metabolic module employable in various pathways is the two-step reaction sequence catalyzed by malate thiokinase (MtkAB) and malyl-CoA lyase (Mcl) (**Fig. 4B, C**). The joint activity of these enzymes activates malate to malyl-CoA (at the expense of ATP) and then cleaves it to generate acetyl-CoA and glyoxylate. This module is part of natural metabolic routes, including natural and modified variants of the serine cycle (Yu & Liao, 2018), but is also required for the HydrOxyPropionyl-CoA/Acrylyl-CoA (HOPAC) cycle (McLean et al., 2023), a synthetic route for CO_2_ fixation so far demonstrated only *in vitro* (**Fig. 4B**). Moreover, this module here has been proposed in the context of the reverse glyoxylate shunt (Mainguet et al., 2013), which in theory allows generation of two C2 compounds from one C4 moiety, and therefore holds potential for achieving higher product yields for C2-dependent productions from sugars (22). Therefore, we reasoned that screening the combined MtkAB and Mcl activity using our metabolic sensors could create new opportunities for pathway engineering. In fact, while the module had previously been shown to rescue an acetyl-CoA auxotrophic strain (Yu et al., 2018), engineering of the full HOPAC cycle requires functionality in a glyoxylate auxotroph because the cycle product is glyoxylate. Hence, we chose to demonstrate the feasibility of the module in our sensor strains as a proof-of-principle for glyoxylate selections.

**Figure 4.**
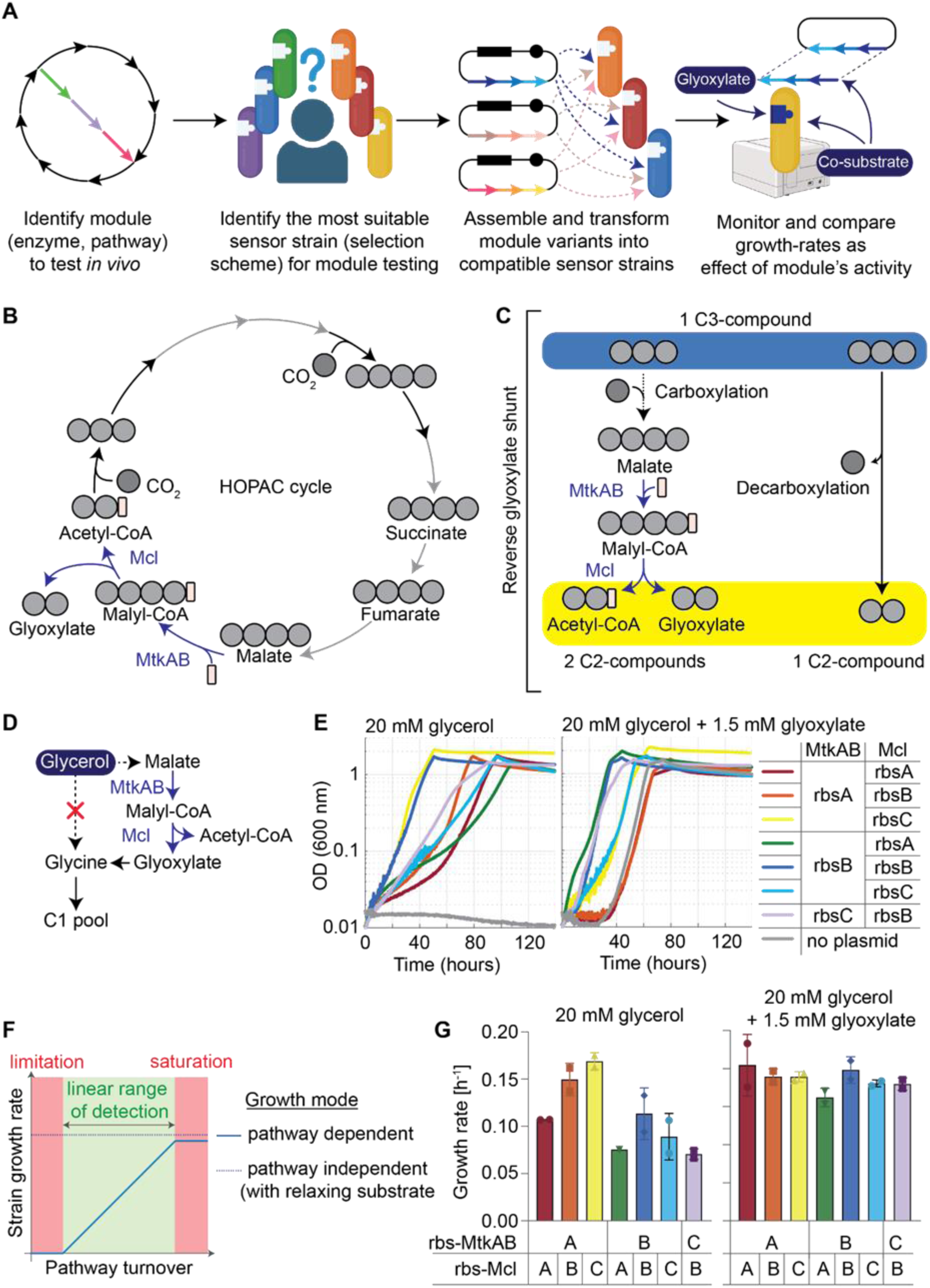
Application of the metabolic sensors for *in vivo* enzyme screeningoperations. (**A**) Workflow for using metabolic sensors with different designs and demands for the screening of metabolic modules. (**B, C**) Lumped architectures of the HydrOxyPropionyl-CoA/Acrylyl-CoA (HOPAC) cycle and reverse glyoxylate shunt. (**D**) Selection scheme of C1+GLY-AUX sensor strain, including MtkAB and Mcl activities. (**E**) Growth profile of the RBS library tested under selective (20 mM glycerol) and non-selective (20 mM glycerol + 1.5 mM glyoxylate) conditions within the C1+GLY-AUX strain. In this type of application for the sensor strain, the growth rate is the proxy for the module activity (metabolic flux). (**F**) Schematic presenting the expected correlation between pathway turnover and growth rate of the selection strain. As long as the module output is too low to allow sensor growth, the growth rate cannot correlate with module activity (“limitation”). On the other hand, if the module turnover is faster than glyoxylate is metabolized, the growth rate will not improve further (“saturation”). In between these extremes, a direct correlation of module turnover and growth rate is expected. (**G**) Quantification of the growth rates for the different candidates under selective and non-selective conditions. Created with Biorender.com.

Growth in the metabolic sensors is expected to be a trade-off between flux capacity through the target module and the metabolic burden associated with its production. Using the growth rate of the metabolic sensor as a proxy for this trade-off, we proceeded by testing how the module capacity is affected by modulating the expression of the required genes (*mtkAB* and *mcl*). We hypothesized that if modules display different flux capacities, they might differ in their growth rate, yet they should not differ in their final biomass density because they have access to the same nutrients in the cultivation medium.

We cloned *mtkAB* and *mcl* under the transcriptional control of RBSs, RBS^A^, RBS^B^ and RBS^C^ as strong, medium and weak ribosome binding sites, respectively (Wenk et al., 2018). Then, we transformed this library into three metabolic sensors: C1+GLY-AUX (**Fig. 4D, E**), 2OXO-AUX (**Supplementary Fig. 10A-D**), and UPP-AUX (**Supplementary Fig. 9E, F**). All C1+GLY-AUX strains grew to the same final optical density (**Fig. 4E**) in line with our above-mentioned expectation that the growth rate would correlate with module flux, while the final optical density should be comparable (**Fig. 4F**). Indeed, different expression constructs resulted in different growth rates, with two constructs resulting in faster growth as a reflection of the combined effect of high flux through MtkAB and Mcl and low metabolic burden (**Fig. 4G**). Therefore, requirement of a strong RBS^A^ for *mtkAB* and weaker RBSs (RBS^B^ or RBS^C^) for *mcl* suggests that the rate-limiting step of the module is the step catalyzed by MtkAB. In summary, we conclude that the following constructs – RBS^A^·*mtkAB-*RBS^B^·*mcl* and RBS^A^·*mtkAB-*RBS^C^-*mcl* – would permit the best trade-off (and therefore the highest flux capacity to support growth) through the module.

Notably, the two other tested selection strains were not suitable for such turnover estimations. The 2OXO-AUX strains showed differences in the final biomass density as well as in growth rates, and required supplementation with three times more succinate when MtkAB-Mcl was expressed (**Supplementary Fig. 10B**). The different behavior of 2OXO-AUX could be a consequence of the selection scheme architecture. With C1+GLY-AUX the fate of the carbon provided by the substrate (glycerol) follows a linear path all the way to glyoxylate (**Fig. 4D**), whereas in 2OXO-AUX succinate is needed both for production of glyoxylate as well as for its further condensation to form isocitrate (**Supplementary Fig. 10C**). Therefore, MtkAB-Mcl and isocitrate lyase may be competing for the succinate pool, which affects the pool of 2-oxoglutarate and therefore the ability to rescue strain growth (**Supplementary Fig. 10A-D**). The last design tested (UPP-AUX) did not result in growth of any of the combinations screened (**Supplementary Fig. 10E, F**). This suggests that the demand from the strain is too high to be supported by the module through its own turnover rate (**Supplementary Fig. 10F**), which implies that current module flux is incapable of supporting growth when the GBR is 0.13 or higher.

We thus demonstrated that different translation rates of MtkAB and Mcl affect the growth rates of the metabolic sensors. These results can be used as a basis to further investigate the corresponding fluxes for glyoxylate production *in vivo*. In the next section, to demonstrate a second application of the metabolic sensors, we focused on quantifying extracellular glycolate in the spent medium of microbial cultivations.

### Detection of glycolate produced through photorespiration

Another aim of this study was to demonstrate the feasible use of metabolic sensors to monitor phenomena of ecological relevance. From an environmental perspective, glycolate synthesis occurs during phytoplankton blooms in the ocean (Lau et al., 2007). During this event, glycolate synthesis is caused by a metabolic process called photorespiration, which is known to limit CO_2_ fixation via the Calvin-Benson-Bassham (CBB) cycle in phototrophic organisms (Bauwe et al., 2010; Dellero et al., 2016). Here, photorespiration occurs through the promiscuous oxygenase activity of the enzyme Ribulose-1,5-bisphosphate carboxylase/oxygenase (RuBisCO) and limits the CO_2_ fixation rate of this enzyme (Federici et al., 2023; Zhu et al., 2010).

Photorespiration has been associated with the secretion of glycolate in photoautotrophic microorganisms in the range of µg/L (< 1 mM, in open waters) to g/L (> 30 mM, under controlled conditions) (Taubert et al., 2019; Yang et al., 2021). Therefore, we reasoned that the sensitivity range of our sensor strains should be able to detect glycolate in the spent medium of phototroph cultures (**Fig. 5A**). Importantly, for this purpose, we reasoned that presence of glycolate will restore growth of our metabolic sensors and that the final optical density will be proportional to the concentration of glycolate in the media.

**Figure 5.**
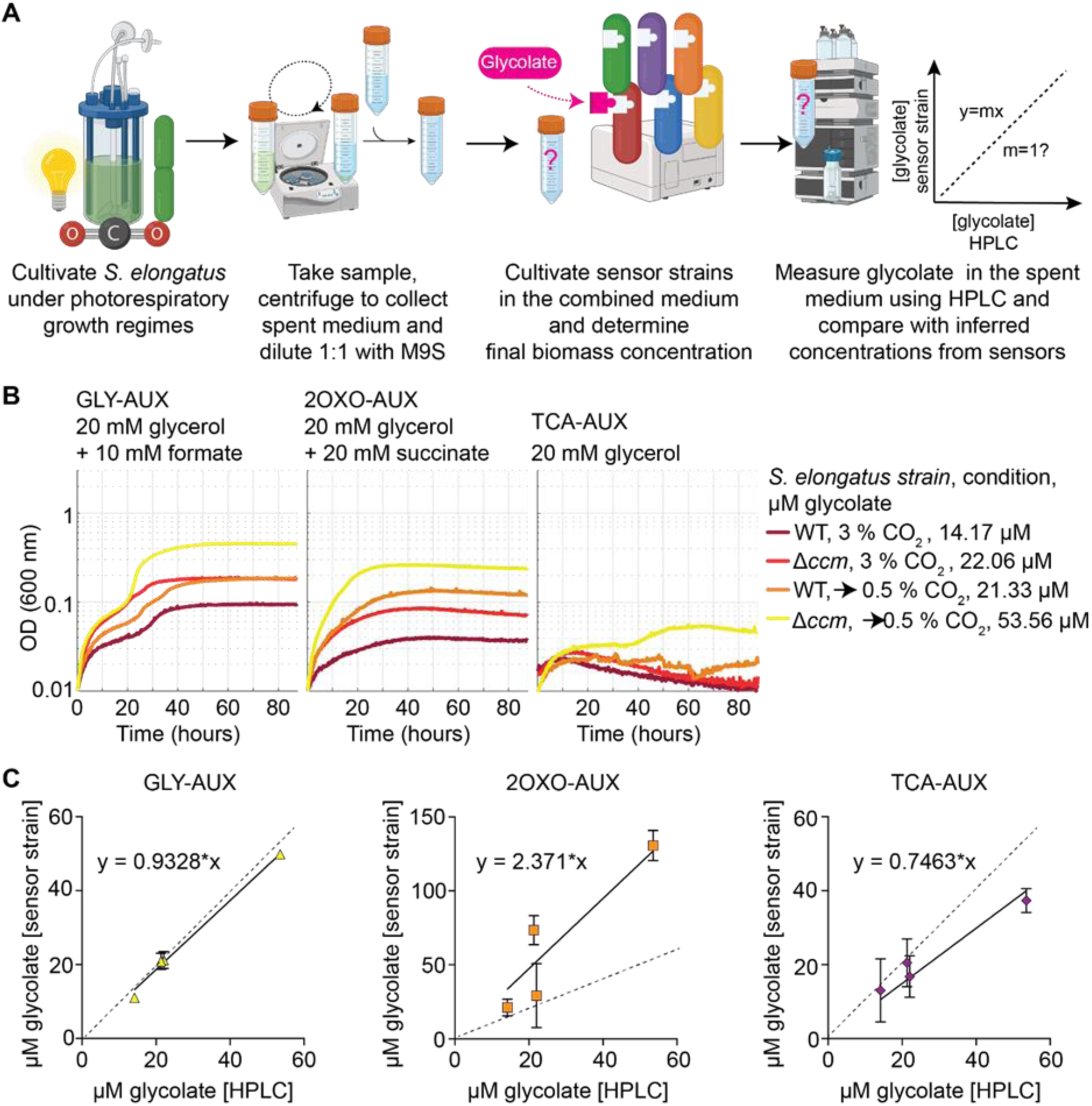
Application of the metabolic sensors for the determination of an extracellular metabolite (glycolate) in the spent medium of a cultivation (photoautotrophic growth). (**A**) Workflow depicting the steps required for the use of metabolic sensors for the determination of the extracellular metabolite in the spent medium of phototrophic *Synechococcus elongatus* strains grown under photorespiratory regimes. (**B**) Growth profile of three metabolic sensors (GLY-AUX, 2OXO-AUX, and TCA-AUX) on the spent medium for the *S. elongatus* strains. The measured glycolate concentrations are shown for each of the conditions tested (cultivation at a constant 3 % CO_2_, or transition from 3 % to 0.5 % CO_2_ to induce photorespiration in a wild type (WT), or carboxysome mutant (Δ*ccm*) background. (**C**) Correlation between the data obtained from HPLC analyses (x-axis) and the concentrations determined by observing the final biomass concentration of different metabolic sensors (y-axis). A good correlation should correspond to a slope close to 1. Created with Biorender.com.

We considered two phototrophic organisms where glycolate secretion was already demonstrated: *Synechococcus elongatus* (Yang et al., 2021) (which grows in fresh water) and *Chlamydomonas reinhardtii* (Taubert et al., 2019) (which grows in salt water). We observed that only the low-salt medium used for *S. elongatus* cultivation was compatible with our *E. coli* strains (**Supplementary Fig. 11 A, B**), and therefore we further investigated glycolate secretion by this cyanobacterium using both wild-type and Δ*ccm S. elongatus* strains. The Δ*ccm* strain lacks the carboxysome, a microcompartment that permits increased local CO_2_ concentrations around RuBisCO and enhances the enzyme’s carboxylation rate (therefore the Δ*ccm* strain is known to exhibit higher rates of RuBisCO oxygenation in ambient CO_2_ conditions (Kaplan & Reinhold, 1999)).

In the experimental setup, we performed cultivation as follows: i) high CO_2_ concentrations (3%) for three days (as negative control where photorespiration should be minimal); ii) alternatively, we cultivated the strains at 3% CO_2_ for three days to reach high biomass densities, followed by an additional three days at 0.5% CO_2_ to stimulate photorespiration. We collected samples at the end of these cultivations, removed cyanobacterial cells by centrifugation, and mixed spent medium in a 1:1 ratio with fresh M9 medium for application of the different metabolic sensors. We then monitored growth of the *E. coli* strains until the stationary phase was reached (**Fig. 5B**) and used the final optical density (adjusted to a 1:1 dilution) to infer glycolate concentration from glyoxylate-to-OD_600_ correlation as discussed above (**Fig. 3D**). As the control, we also measured the glycolate concentrations in the spent media samples using conventional analytics (HPLC). As expected, the Δ*ccm* strain grown under photorespiratory conditions yielded the highest glycolate concentration, whereas the wild-type strain grown constantly at high CO_2_ resulted in the lowest concentration (**Fig. 5B**). This result was confirmed by the final optical density values in all three sensor strains, which thus suggests that optical density measurements can serve as at least a coarse estimate of the relative glycolate concentration within the samples.

We continued by determining which of the three strains was most precise in sample detection. From the HPLC data, we could confirm that the concentration of the samples ranged from 10 µM to 50 µM glycolate (**Fig. 5B**), which is within the linear range of our most sensitive GLY-AUX strain (**Fig. 3D**). In fact, when plotting the sensor-derived concentration estimates of glycolate against those measured with HPLC, we could determine a slope of 0.93 for the GLY-AUX measurements (R^2^ = 0.992) (**Fig. 5C**). On the other hand, since the other two strains 2OXO-AUX and TCA-AUX presented a linear range for glyoxylate-to-OD_600_ correlation only at higher glycolate concentrations (**Fig. 3D**), we observed a greater discrepancy between HPLC-derived and sensor-derived measurements for these strains, which resulted in a slope of 2.37 for 2OXO-AUX and 0.75 for TCA-AUX. Therefore, when glycolate concentrations lay within the linear range of detection of the sensor, measurement of the strain’s final optical density can be a reliable readout of the concentration of the real target molecule in the medium.

## Discussion

Metabolic sensors are valuable tools for several biological applications, including synthetic biology and environmental monitoring. Yet their implementation is not trivial because the created auxotrophic phenotypes must be robust in various contexts. Achieving such phenotypes requires deep knowledge of the host’s metabolism. In our study, we streamlined the workflow for creating such sensor strains by combining the *in silico* design of metabolic sensors with their engineering and physiological characterization together with *in vivo* applications. We selected glyoxylate as the metabolite-of-interest around which we created a set of sensor strains because this metabolite is not essential for growth and therefore creating *ad hoc* growth-coupled designs requires rewiring of the host’s metabolism. Through this workflow, we created a set of six sensor strains available off-the-shelf for the same screening purpose. We showcased their application in two practical cases where growth rate or final optical density served as a proxy for prototyping a segment of an engineered metabolic pathway and for environmental monitoring, respectively. Having access to multiple strains allowed us to successfully accomplish both these screening investigations.

To implement the sensor construction workflow, we used a medium-scale metabolic model (Corrao et al., 2024) to predict enzyme inactivations which force growth to be dependent on glyoxylate. The procedure for finding suitable deletion candidates had been previously applied to design strains that depend on CO_2_-fixation via RuBisCO (N. Antonovsky et al., 2016), glycerate sensor strains (Aslan et al., 2020), or synthetic methylotrophs (Keller et al., 2020; Keller et al., 2022). Yet, in all these cases, computational analysis relied on a metabolic core model (Orth et al., 2010) which covered only glycolysis/gluconeogenesis, the pentose phosphate pathway, the TCA cycle, and oxidative phosphorylation. In contrast, the larger size of the model used in this study allowed us to explore more designs than with the core *E. coli* model while reducing computation time and unfeasible solutions typical of genome-scale models (Monk et al., 2013). Thanks to the algorithm, we could identify three groups of design strategies (glycolytic disruptions, interruption of TCA anaplerosis, and transamination of glyoxylate to glycine) which we further engineered in the sensor strains. In particular, the glyoxylate to glycine transamination group of solutions was possible thanks to this medium-scale model because it relied on enzyme inactivations within the amino acid biosynthetic network.

To fully implement robust metabolic sensors, we had to complement the list of candidate enzymes proposed by the algorithm with additional gene deletions. Most of these interventions were inspired by data available in databases such as EcoCyc (https://ecocyc.org/) (Karp et al., 2023). We applied the following types of manual interventions to complement the model: deletion of latent metabolic pathways which may become activated, e.g. Δ*mgsA* (Iacometti et al., 2022; Long & Antoniewicz, 2019b) in LOW-AUX; addition of heterologous modules to relieve metabolic demands, i.e., C1 moieties, as in the case of the C1 module from the reductive glycine pathway (Yishai et al., 2017) in GLY-AUX; limitation in reaction annotations from a thermodynamic perspective, as in the case of *maeAB*, since ME1 and ME2 were considered irreversible in metabolic models; limitation in annotations as an effect of promiscuous reactions, such as in the case of *ltaE* (in C1+GLY-AUX and GLY-AUX) and *prpC* (in 2OXO-AUX), which can complement inactivation of *glyA* (Liu et al., 1998) and *gltA* (Gerike et al., 1998), respectively. Ultimately, *in vivo* characterization of the designed strains confirmed the model predictions in terms of the ranked sensitivity of the strains towards glyoxylate (and glycolate).

The main application of metabolic sensors is for testing enzyme modules by measuring growth as a proxy for their activity (Orsi et al., 2021; Wenk et al., 2018), where growth reflects a trade-off between the module’s flux capacity and the metabolic burden for its synthesis. The two-carbon molecule glyoxylate is a product of CO_2_ fixation pathways, such as the serine cycle or the *in vitro* crotonyl-CoA/ethylmalonyl-CoA/hydroxybutyryl-CoA (CETCH) cycle and the HOPAC cycle (McLean et al., 2023; Schulz-Mirbach et al., 2024), and can be synthesized through the “reverse glyoxylate shunt” from pyruvate bypassing its decarboxylation to acetyl-CoA (Mainguet et al., 2013; Orsi et al., 2022). We tested MtkAB and Mcl as key components of the reverse glyoxylate shunt and HOPAC cycle (Mainguet et al., 2013; McLean et al., 2023), with indications that stronger expression of MtkAB is required compared to Mcl. Experimental data from testing different modules suggested that choice of the selection scheme (and therefore metabolic sensor) also has an impact on the ability to screen for modules. We demonstrated that strains with high metabolic demands or competitive nodes for substrate utilization might not be suitable for module testing.

Furthermore, we demonstrated how the estimation of extracellular metabolites (glycolate) in spent cultivation medium is most precise when using metabolic sensors whose linear range of detection overlapped with the target molecule concentration in the medium. In the literature, a similar approach was used to determine mevalonate in spent medium but that approach used growth rate as the benchmark (Pfleger et al., 2007). Instead, here we demonstrated how the final biomass concentration can also be used as a proxy for determining the concentration of the target molecule in the spent medium, when interpolated with the compound-to-OD_600_ correlation. The glycolate metabolic sensors we created exhibited a linear range of detection within the concentration range 0.01-20 mM, thus adding to a list of transcription-based [0.1-200 mM range (Xu et al., 2021) or 0.01-20 mM range (Barthel et al., 2024)] and enzyme-based [0.01-1 mM range (Tsiafoulis et al., 2002)] biosensors previously described in the literature. We argue that this simple setup for measuring the molecule of interest can be of use when performing high-throughput screening using multi-well-plate setups.

We postulate that using a computational algorithm complemented by manual interventions will streamline the creation of growth-coupled designs for various metabolites of interest. The metabolic sensors generated in the presented study will allow us to explore novel designs for synthetic pathways leading to the synthesis of glyoxylate. Moreover, the auxotrophic phenotype can be exploited in this way for the evolution of (new) key enzymatic steps leading to synthesis of the target metabolite (Chen & Arnold, 2020; McLure et al., 2022; Molina et al., 2022; Wang et al., 2021) as well as in high-throughput automated setups (Li et al., 2023; Orsi et al., 2024; Yu et al., 2023). Since glyoxylate is described as a key intermediate of some protometabolic pathways (Krishnamurthy & Liotta, 2023; Pulletikurti et al., 2022; Springsteen et al., 2018; Yadav et al., 2022), we believe our sensor strains could create new possibilities for explorative studies within this field of research. Finally, with glycolate as a marker for photorespiration, we encourage the use of such metabolic sensors for high-throughput study of photorespiration by other organisms, which has broad implications for agriculture (Betti et al., 2016; Garcia et al., 2023; Walker et al., 2016) and environmental studies (Bauwe et al., 2010; Behrenfeld & Boss, 2014).

## Materials and Methods

### The stoichiometric model

In order to identify promising knockout strategies in *E. coli* that create glyoxylate auxotrophy at different dependency levels, we used the recently published iCH360 model (Corrao et al., 2024). iCH360 is a subnetwork of the much larger genome-scale model, focusing on central metabolic subsystems that carry relatively high flux, are central to maintaining and reproducing the cell, and provide precursors and energy to engineered metabolic pathways. This medium-sized model, by doing away with low-flux and secondary pathways and enzymes, facilitates applications like ours since these redundant reactions typically create bypasses that are unlikely to be relevant *in vivo* and greatly complicate the search. However, since we wanted to design auxotrophic strains to a non-standard carbon source (glyoxylate), we had to augment the iCH360 model with 2 metabolites and 5-6 reactions in order for it to be able to deal with glyoxylate metabolism:

#### Metabolites

- extracellular glyoxylate (glx_e): formula = C3H3O4
- 2-Hydroxy-3-oxopropanoate (2h3oppan_c): formula = C2H1O3 Note that 2-Hydroxy-3-oxopropanoate and tartronate semialdehyde are synonyms.

#### Reactions

- glyoxylate exchange (EX_glx_e): glx_e ⇔
- glyoxylate transport (glx_t): glx_e + h_c ⇔ glx_c + h_c
- aspartate-glyoxylate transaminase (BHC): asp L_c + glx_c ⇔ oaa_c + gly_c
- glyoxalate carboligase (GLXCL): glx_c + h_c ⇔ 2h3oppan_c + co2_c
- tartronate semialdehyde reductase (TRSARr): 2h3oppan_c + h_c + nadh_c ⇔ glyc R_c + nad_c
- formate-tetrahydrofolate ligase (FTHFLi*): for_c + atp_c + thf_c ⇔ 10fthf_c + adp_c + pi_c

* The FTHFLi reaction was added only in the strain named C1+GLYAUX which utilizes formate as the source for all the C1 carbon metabolism.

### Systematic search for growth-coupled designs

After establishing a suitable stoichiometric model as described in the previous section, we applied a search algorithm for identifying auxotrophic knockout strategies.

First, ATP maintenance reaction, as the algorithm is designed to only calculate the marginal dependence on glyoxylate (at low growth rates) and the maintenance reaction is not relevant for that calculation. Then, we set the bounds of the glucose exchange flux to 0 (instead of the default lower bound of -10 mmol/gCDW/h), and replaced it with succinate or glycerol as the abundant carbon source, by setting the lower bound to -1000 mmol/gCDW/h.

The next step is to iterate through all possible single or double knockouts of reactions from the list of central reactions - i.e. reactions that exist in the core model of *E. coli* (Orth et al., 2010) and the ones added to iCH360 (excluding the glyoxylate uptake itself). In some cases, we lumped together two reactions (denoted as REACTION1|REACTION2), either because they are arranged in one linear pathway with no branchpoints, catalyzed by the same enzyme, or the same chemical reaction with different cofactors (catalyzed by two isoenzymes):

- **G6PDH2r**
- **PGL**
- **PGI**
- **PFK**
- **FBP**
- **FBA**
- **TPI**
- **PGK**
- **GAPD**
- **PGM**
- **ENO**
- **PYK**
- **PPS**
- **PDH**
- **PFL**
- **GND**
- **RPE**
- **RPI**
- **TKT1|TKT2**
- **TALA**
- **PPC**
- **PPCK**
- **ME1|ME2**
- **SUCDi**
- **FUM**
- **CS**
- **ACONTa|ACONTb**
- **ICDHyr**
- **ICL**
- **MALS**
- **AKGDH**
- **SUCOAS**
- **MDH**
- **ALCD2x**
- **ACALD**
- **GLXCL|TRSARr**
- **GHMT2r**
- **BHC**

For each possible single or double knockout, we ran a series of flux-balance analyses (FBA) using the cobrapy toolbox (Ebrahim et al., 2013). We first tested whether it can grow at all on succinate. If it could, we ran another FBA with a high abundance of succinate, and a limiting amount of glyoxylate. The ratio between the maximal growth rate and the glyoxylate uptake rate represents that GBR. The same is repeated with glycerol instead of succinate. We then normalized the GBR value by dividing them by the GBR of a wild-type cell growing on glyoxylate alone (without succinate nor glycerol). This brings the values to generally be between 0 and 1, except for a few cases where the KO makes the cell require even more glyoxylate (typically, this is not directly related to the glyoxylate itself, but rather a disruption that makes biosynthesis less efficient and therefore requiring more ATP). The results are summarized in **Supplementary Figure 1**. Based on these results, we selected 5 designs that span the range of relevant GBR values (**Supplementary Table 1**).

### Strains and plasmids used in this study

All strains and plasmids employed in this study are listed in **Supplementary Table 2**. *E. coli* SIJ488, which carries recombinases behind an arabinose inducible promoter and a flippase behind a rhamnose dependent promoter (Jensen et al., 2015), was the base strain for all engineering efforts and was used as wildtype reference whenever required. NEB5α cells were used for cloning.

### Gene deletions

Gene deletions were performed by λ-red recombineering or through Cas9-mediated base editing to introduce premature STOP codons (Volke et al., 2022). For λ-red recombineering, knockout cassettes were constructed by amplifying the antibiotic resistance cassette from pKD3 (chloramphenicol resistance, specified by “Cap” in primer name) or pKD4 (kanamycin resistance) (both vectors were a gift from Barry L. Wanner; Addgene plasmids #45604 or #45605) with ‘KO’-primers introducing 50 bp overhangs with homology to the respective target locus using PrimeStar GXL polymerase (Takara Bio). These ‘KO’-primers were designed following the KEIO knockout collection homology arms sequences (Baba et al., 2006). After purifying the cassette by PCR purification using the GeneJet PCR purification kit (Thermo Scientific, Dreieich, Germany), the target strain was electroporated with the fragment. For this, the cells were inoculated in LB medium and grown to an OD_600_ of 0.3 – 0.5, when recombinase expression was induced by adding 15 mM L-arabinose and further incubation for 45 min at 37°C. After harvesting the cells by centrifugation (11,000 rpm, 30 sec, 2°C) and washing them three times with ice cold 10 % glycerol, the cells were electroporated with with ∼300 ng of the deletion cassette (1 mm cuvette, 1.8 kV, 25 µF, 200 Ω). The cells were plated on LB plates containing the relevant antibiotic. Successful gene deletion in grown colonies was confirmed by verifying the target locus size in a PCR using ‘ver’-primers (**Supplementary Table 3**) and DreamTaq polymerase (Thermo Scientific, Dreieich, Germany). To remove the antibiotic resistance cassette, 2 ml strain culture was grown to an OD_600_ of 0.5, when 50 mM L-rhamnose were added to induce flippase expression, followed by incubation for ≥ 3 h at 30 °C. To confirm successful cassette removal, the colonies were tested for antibiotic sensitivity and the target locus size was confirmed by PCR with ‘KO-Ver’ primers and DreamTaq polymerase (Thermo Scientific, Dreieich, Germany).

For Cas9-mediated base editing, we followed the protocol provided by the original work (Volke et al., 2022). We constructed pMBEC plasmids containing up to six single-guide RNAs (sgRNAs) for targeting up to three genes (two sgRNA/gene) per round of mutation. The spacer sequences for premature STOP codon introduction were identified using the CRISPy-web tool (https://crispy.secondarymetabolites.org/, (Blin et al., 2016)) and are listed in **Supplementary Table 4**.

### Gene integration by P1 transduction

To reintroduce *glcDEF* in the TCAAUX strain (Claassens et al., 2020), *glcB* was deleted by P1 transduction (Thomason et al., 2007), thus transferring the *glcB* deletion locus with a kanamycin resistance from a Δ*glcB* donor strain (JW2943) from the KEIO collection (Baba et al., 2006) and the adjacent *glcDEF* wildtype genes. After the transduction, the cells were plated on LB-kanamycin plates containing 20 mM sodium citrate. Successful reintroduction of *glcDEF* was confirmed by PCR using the verification primers previously used to verify the deletion of *glcDEFGB* (**Supplementary Table 2**). Furthermore, growth of the strain with glycolate instead of glyoxylate was confirmed in tubes.

### Plasmid-based gene expression

For plasmid-based gene expression, the pZ-ASS plasmid (p15A origin of replication and strong promoter)was used (Wenk et al., 2018). Malate thiokinase (*mtkAB* from *Methylococcus capsulatus*, as described by Liao et al. (Mainguet et al., 2013)) and the codon-optimized variant of malyl-CoA lyase (*mcl* from *Rhodobacter spaeroides*) was obtained from pTE3262 (unpublished, both plasmids were provided by Dr. Shanshan Luo, for gene sequences see **Supplementary Table 5**). All genes were amplified with primers introducing ribosomal binding sites “A”, “B” or “C” (Zelcbuch et al., 2013) as indicated by the primer name, the coherent overlapping primers were used to amplify the pZ-ASS backbone (Supplementary Table 2). In front of all genes, a spacer of the sequence “TAATAGAAATAATTTTGTTTAACTTTA” was introduced. In addition, between *mtkB* and *mtkA* another spacer of the sequence “TCTAGAGCTAGCGTTGATCGGAGGTTCTGTTAAGTAACTGAACCC” was introduced, while between *mtkA* and *mcl* a spacer of the sequence “TGTCGTTAGTGACGCTTACCTCTT” was introduced to achieve non-redundant fragment overhangs. The fragments needed to obtain pZ-ASS-MtkBA-Mcl plasmids with combinations or different ribosomal binding sites were assembled by a HiFi DNA assembly as described for the HiFi assembly protocol (New England Biolabs, Ipswich, Massachusetts). After transforming the HiFi NEB5α cells with the plasmid, the cells were plated on LB Streptomycin plates. Successful plasmid assembly was confirmed by whole-plasmid sequencing (plasmidsaurus, Eugene, Oregon).

### Routine strain cultivation

For routine strain handling, Lysogeny broth (LB) medium (composed of 1% NaCl, 0.5% yeast extract, 1% tryptone) was used. When appropriate, antibiotics (kanamycin (25 μg/mL), ampicillin (100 μg/mL), streptomycin, (100 μg/mL), or chloramphenicol (30 μg/mL)) were added.

### Plate reader experiments

For growth tests, M9 minimal medium without antibiotics was used (50 mM Na_2_HPO_4_, 20 mM KH_2_PO_4_, 20 mM NH_4_Cl, 2 mM MgSO_4_, 1 mM NaCl, 134 μM EDTA, 100 μM CaCl_2_, 13 μM FeCl_3_·6H_2_O, 6.2 μM ZnCl_2_, 1.62 μM H_3_BO_3_, 0.76 μM CuCl_2_·2H_2_O, 0.42 μM CoCl_2_·2H_2_O, 0.081 μM MnCl_2_·4H_2_O). Carbon sources were added as indicated in the text. For “relaxing” conditions of overnight precultures, 1.5 mM glyoxylate or glycolate were supplemented, depending on the experiment. For the experiment, 2 ml culture were harvested by centrifugation (10,000 g, 30 sec) and washed three times with “selective” M9 minimal medium in the absence of glyoxylate or glycolate. Then, the washed cells were diluted to a final OD_600_ of 0.01 in a 96-well microtiter plates (Nunclon Delta Surface, Thermo Scientific). The medium composition (relaxing, selective, or relaxing with dilutions of glyoxylate or glycolate) was adjusted for each experiment. Each well of the 96 well plate contained 150 μL of culture and 50 μL mineral oil (Sigma-Aldrich) to prevent evaporation but allow gas exchange. A BioTek Epoch 2 plate reader (BioTek, Bad Friedrichshall, Germany) was used to monitor growth of technical duplicates at 37 °C by measuring the absorbance (at 600 nm) every ∼10 minutes with intermittent orbital and linear shaking. During the analysis with MATLAB, blank measurements were subtracted and OD_600_ values were converted to cuvette OD_600_ values by multiplying with a factor of 4.35 which had previously been established for the instrument.

### 13C labelling experiments

For stationary isotope labelling of proteinogenic amino acids ^13^C_2_-glyoxylate ([1,2-^13^C]glyoxylic acid monohydrate, LGC Standards) was used as tracer. As control, sodium glyoxylate monohydrate (Sigma Aldrich) was used as unlabelled substrate. All experiments were performed in triplicate as follows. Strains were grown in 4 mL LB. Subsequently, 40 µL of culture was transferred into 4 mL of fresh M9 medium supplemented with labelled or unlabelled tracer plus the additional co-substrate (glycerol or succinate). To reduce the effect of LB carryover, once the OD_600_ reached a level of 0.8, 40 µL of grown culture were transferred again into 4 mL of the same M9 medium and at late exponential phase, the equivalent of 1 mL of culture at OD_600_ of 1 was harvested. Samples were then processed to analyse proteinogenic amino acid mass-isotopomers through GC-MS (Donati et al., 2023). For analysis of amino acid isotopomer data, we considered fragments containing the full carbon backbone of the amino acids of interest (Long & Antoniewicz, 2019a). Raw data from the GCMS was integrated using SmartPeak (Kutuzova et al., 2020). Processed data was further corrected for the natural abundance of isotopes in the derivatization agents used for GCMS analysis (Wahl et al., 2004).

### Proteomics

The mass spectrometry proteomics data have been deposited to the ProteomeXchange Consortium via the PRIDE partner repository with the dataset identifier PXD054993. C1+GLY-AUX and a wild type were cultivated in 10 ml of M9 + 20 mM glycerol + 1.5 mM glyoxylate or M9 + 20 mM glycerol + 5 mM glycine at 37 °C shaking at 180 rpm. Cells from three biological replicates were harvested at an equivalent of 1 ml of OD 3 in the late exponential phase by centrifugation for 1 minute at 11,000 g. The cells were washed two times with phosphate buffer (12 mM phosphate buffer, 2.7 mM KCl, 137 mM NaCl, pH=7.4) before being flash frozen in liquid nitrogen for subsequent storage at -70 °C. To isolate the proteome, cell pellets were resuspended in 2% sodium lauroyl sarcosinate (SLS, in 100 mM ammonium bicarbonate) and heat incubated at 95°C for 15 min. For DNA shearing, the cells were then sonicated for 30 seconds (Vial Tweeter, Hielscher). After quantifying the total protein in a BCA assay, the samples were treated with 5 mM Tris (2-carboxy-ethyl) phosphine (TCEP) at 90°C for 15 min, followed by protein alkylation using 10 mM Iodoacetamide for 30 minutes in the dark at 25 °C. For protein digestion, 50 µg of total protein were incubated with 1 µg of porcine trypsin (Promega) in presence of 0.5 % SLS at 30 °C overnight. After digestion, SLS was precipitated by adding a final concentration of 1.5% trifluoroacetic acid (TFA, Thermo Fischer Scientific). Peptides were desalted by using C18 solid phase extraction cartridges (Macherey-Nagel). Cartridges were prepared by adding acetonitrile (ACN), followed by equilibration with 0.1% TFA. Peptides were loaded on equilibrated cartridges, washed with 5% ACN and 0.1% TFA containing buffer, eluted with 50% ACN and 0.1% TFA and finally dried. The peptides were resuspended in 100 µl 0.1% TFA and the peptide mixtures were then analyzed by LC-MS on an Exploris 480 instrument connected to an Ultimate 3000 RSLC nano and a nanospray flex ion source (Thermo Scientific). A capillary column (75 μm x 42 cm) packed in-house with C18 resin (2.4 μm, Dr. Maisch) was used for peptide separation. The following separating gradients were used: 94% solvent A (0.15% formic acid) and 6% solvent B (99.85% acetonitrile, 0.15% formic acid) to 35% solvent B over 60 minutes at a flow rate of 300 nl/min. DIA-MS acquisition method was performed with: spray voltage set to 2.3 kV, funnel RF level at 45, and heated capillary temperature at 275 °C. For DIA experiments MS1 resolution was set to 120.000 at m/z 200 and full MS AGC target was 300% with an max. injection time (IT) of 50 ms. Mass range was set to 350– 1400. AGC target value for fragment spectra was set at 3000%. 49 windows of 15 Da were used with an overlap of 1 Da. Resolution was set to 15.000 and IT to 22 ms. Stepped HCD collision energy of 25, 27.5, 30 % was used. MS1 data was acquired in profile, MS2 DIA data in centroid mode.

Analysis of DIA data was performed using DIA-NN version 1.8 (Demichev et al., 2020), using the UniProt protein database from Escherichia coli K12 and added sequences for the SIJ488 λ-red recombineering machinery (Phage recombinase gam (UniProt ID NP_040618.1), Phage recombinase beta (UniProt ID WP_000100844), Phage recombinase exo (UniProt ID 1AVQ_A), Flippase (UniProt ID P03870.1). Full tryptic digest was allowed with three missed cleavage sites, and oxidized methionines and carbamidomethylated cysteines. Match between runs and remove likely interferences were enabled. The neural network classifier was set to the single-pass mode, and protein inference was based on genes. Quantification strategy was set to any LC (high accuracy). Cross-run normalization was set to RT-dependent. Library generation was set to smart profiling. DIA-NN outputs were further evaluated using a SafeQuant version modified to process DIA-NN outputs (Ahrné et al., 2013).

### Determining extracellular glycolate in spent medium using LC-MS/MS

Chromatographic metabolite separation was performed on an Agilent Infinity II 1290 HPLC system using a Kinetex EVO C18 column (150 × 2.1 mm, 3 μm particle size, 100 Å pore size, Phenomenex) connected to a guard column of similar specificity (20 × 2.1 mm, 3 μm particle size, Phenomenex) at a constant flow rate of 0.1 mL/min with 2 µl injection volume and 0.1 % formic acid in water as mobile phase A and 0.1 % formic acid in methanol (Honeywell, Morristown, New Jersey, USA) as phase at 25 °C. The mobile phase profile consisted of the following steps and linear gradients: 0 – 4 min constant at 0 % B; 4 – 6 min from 0 % to 100% B; 6 – 7 min constant at 100 % B; 7 – 7.1 min from 100 % to 0% B; 7.1 – 12 min constant at 0 % B. An Agilent 6495 mass spectrometer was used in negative mode with an electrospray ionization source and the following conditions: ESI spray voltage 2000 V, nozzle voltage 500 V, sheath gas 300 °C at 11 L/min, nebulizer pressure 50 psig and drying gas 80 °C at 16 L/min.

Compounds were identified quantified based on their mass transition, retention time and peak ara compared to an external standard curve using the MassHunter software (Agilent, Santa Clara, CA, USA). The parameters mass transitions, collision energies, Cell accelerator voltages, and Dwell times have been optimized using chemically pure standards and are given in **Supplementary Table 6**.

## Acknowledgments

The authors thank Jörg Kahnt for technical assistance with proteomics and Beau Dronsella for valuable feedback on the strain designs.

## Author contributions

EO and HSM conceptualized the study. EN and HH predicted knockout combinations and selection demands in silico. CC and EO identified additional deletion targets with input from SNL and ABE. EO, HSM., CC, AS, HP, SA, NG, RV, FS and AM constructed the strains used in this study. EO, HSM and CC characterized the sensor strains. HSM generated datapoints for the quantitative sensitivity comparisons. EO and HSM assembled results from all authors and wrote the original manuscript draft with help from EN, HH and PIN. EO generated labelling samples, which were processed and analyzed by KSJ and SD. NP quantified metabolites and analyzed the data. AK cultivated S. elongatus strains and TC cultivated C. reinhardtii strains. TG analyzed proteomic samples. EO, ABE, TE and PIN acquired funding. ABE, TE and PIN supervised the research.

## Data and materials availability

All scripts, data files, and result plots can be found on our GitLab repository: https://gitlab.com/elad.noor/glyoxylate-auxotrophy.

