## Supplementary Material for "Expanding the biotechnological scope of metabolic sensors through computation-aided designs"

**This PDF file includes:**

Figs. S1 to S11
Tables S1 to S6

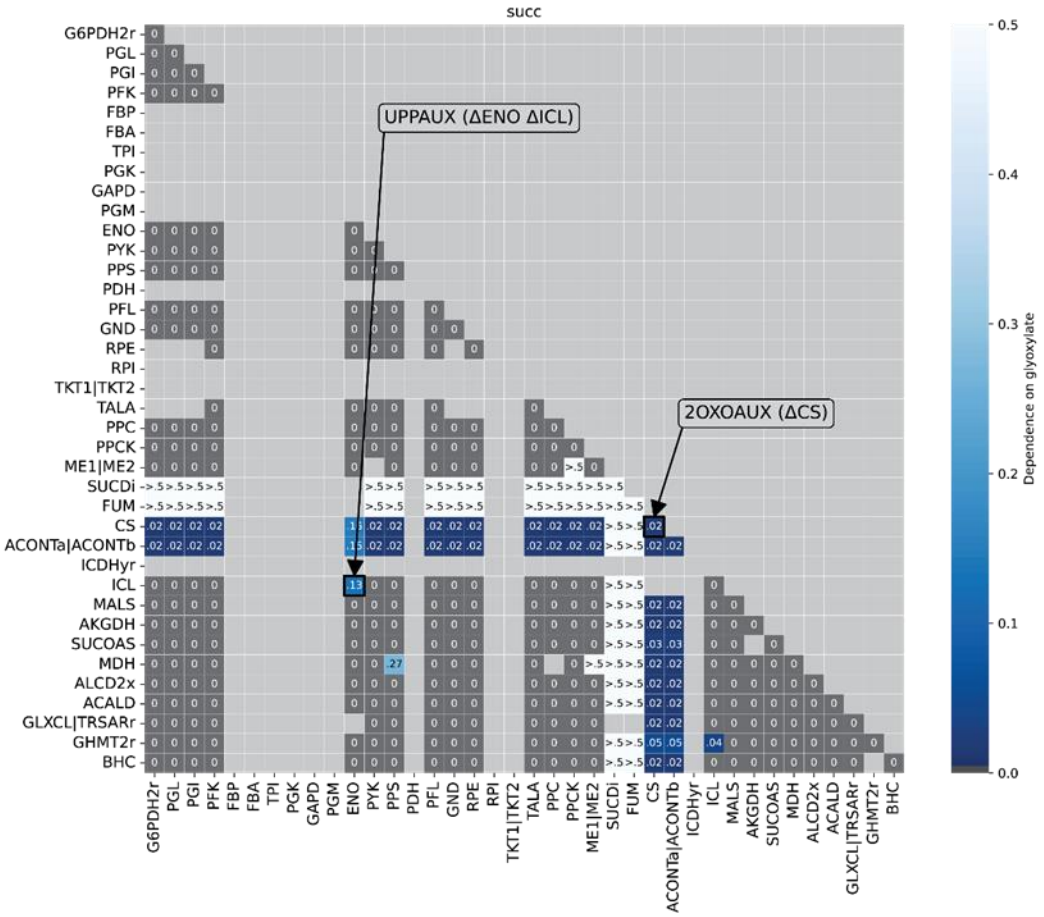
succ
UPPAUX (ΔENO ΔICL)
2OXOAUX (ΔCS)
Dependence on glyoxylate

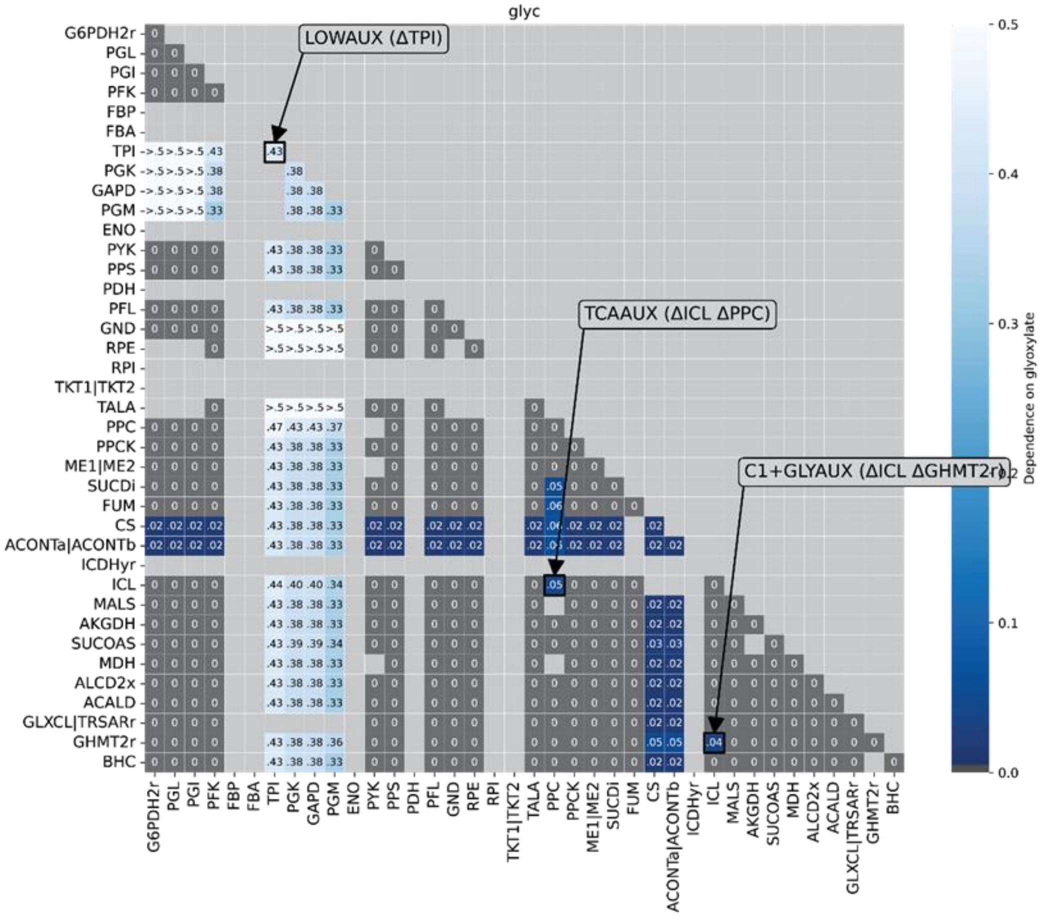
glyc
LOWAUX (ΔTPI)
TCAAUX (ΔICL ΔPPC)
C1+GLYAUX (ΔICL ΔGHMT2r)
Dependence on glyoxylate

24

**Fig. S1.**

Output data from the systematic search for deletions determining an auxotroph phenotype with regard to glyoxylate. Dependency on glyoxylate (represented by normalized GBR, glyoxylate-biomass ratios) for all single/double knockouts tested. Values above 50% dependence are shown as ">.5", as they are outside the range of interest for our biotechnological application.

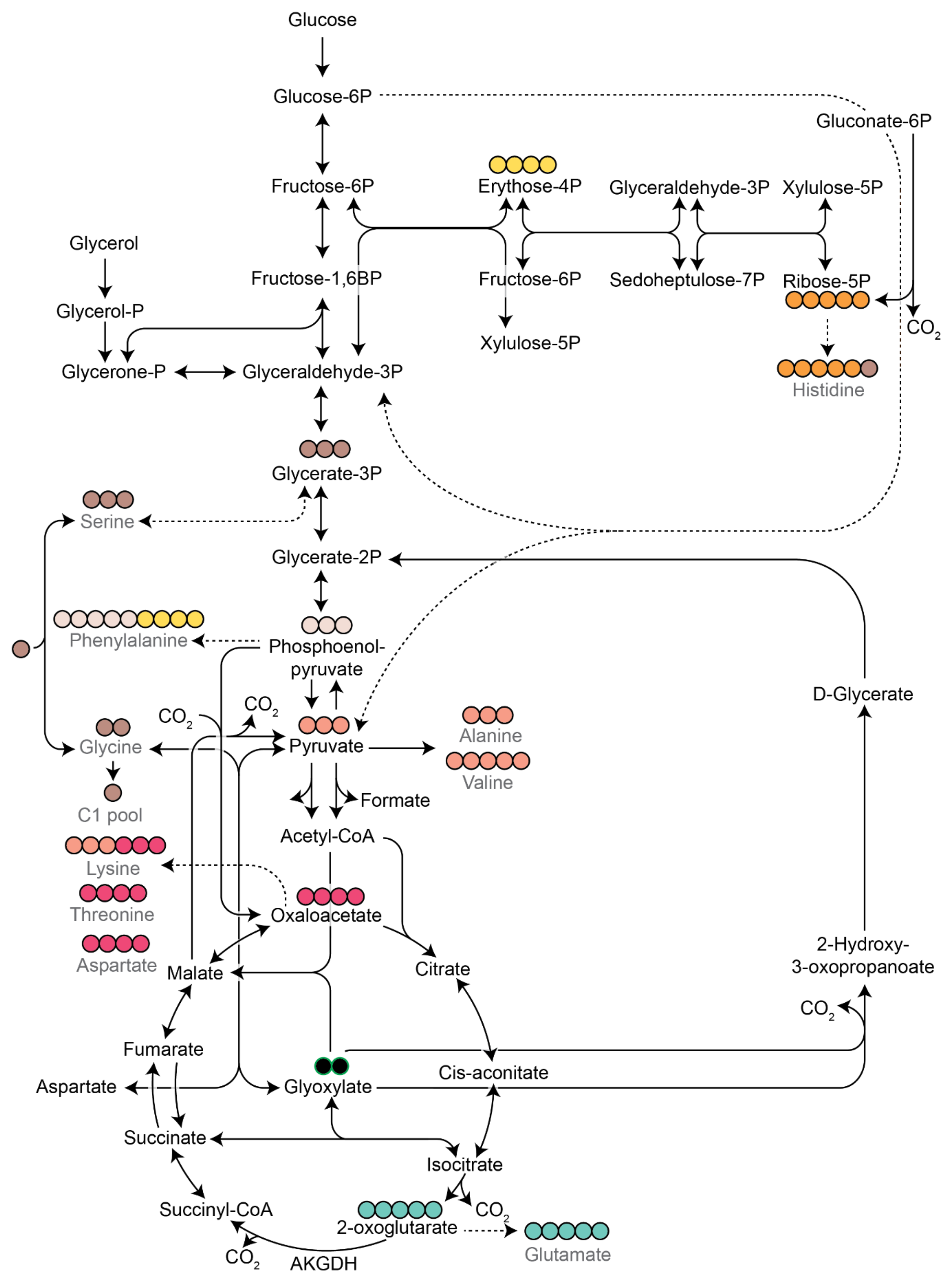
Glucose
Glucose-6P
Gluconate-6P
Fructose-6P
Erythose-4P
Glyceraldehyde-3P
Xylulose-5P
Glycerol
Fructose-1,6BP
Fructose-6P
Sedoheptulose-7P
Ribose-5P
Glycerol-P
$CO_2$
Xylulose-5P
Glycerone-P
Glyceraldehyde-3P
Histidine
Glycerate-3P
Serine
Glycerate-2P
Phenylalanine
Phosphoenol-
pyruvate
D-Glycerate
$CO_2$
$CO_2$
Alanine
Glycine
Pyruvate
Valine
Formate
C1 pool
Acetyl-CoA
Lysine
Threonine
Oxaloacetate
Aspartate
Malate
Citrate
2-Hydroxy-
3-oxopropanoate
$CO_2$
Fumarate
Cis-aconitate
Aspartate
Glyoxylate
Succinate
Isocitrate
Succinyl-CoA
$CO_2$
2-oxoglutarate
Glutamate
$CO_2$
AKGDH

**Fig. S2.**

Origin of amino acids analysed in isotopic labelling experiments in the central metabolism of E. coli. Closed circles represent carbon atoms and are coloured according to the carbon atoms of central metabolism precursors from which they originate from.

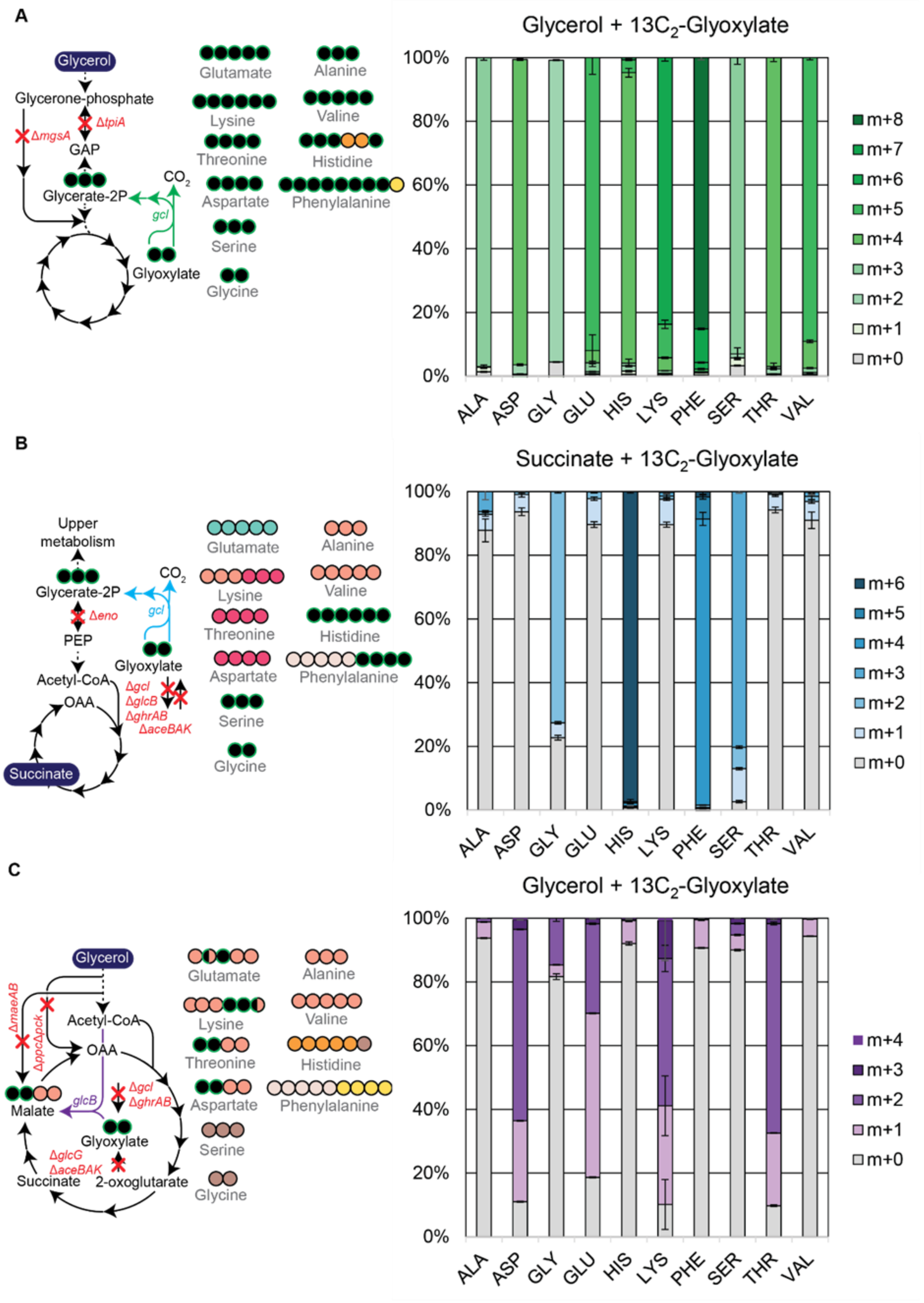


**Fig. S3.**

Labelling patterns for amino acids of metabolic sensor LOW-AUX, UPP-AUX, and TCA-AUX strains grown with $^{13}C_2$ glyoxylate. Circles represent carbon atoms and are coloured according to the carbons from central metabolism precursors they originate from (as shown in Supplementary Figure 2). $^{13}C$ carbon moieties stemming from $^{13}C_2$

glyoxylate are shown as black closed circles with green border. In cases where $^{13}C$ and $^{12}C$ incorporation are equally likely, the respective closed circle is shown in split colors. A) Labelling patterns measured for the LOW-AUX strain grown with glycerol and $^{13}C_2$ glyoxylate. B) Labelling patterns measured for the UPP-AUX strain grown with succinate and $^{13}C_2$ glyoxylate. C) Labelling patterns expected for the TCA-AUX strain grown with glycerol and $^{13}C_2$ glyoxylate.

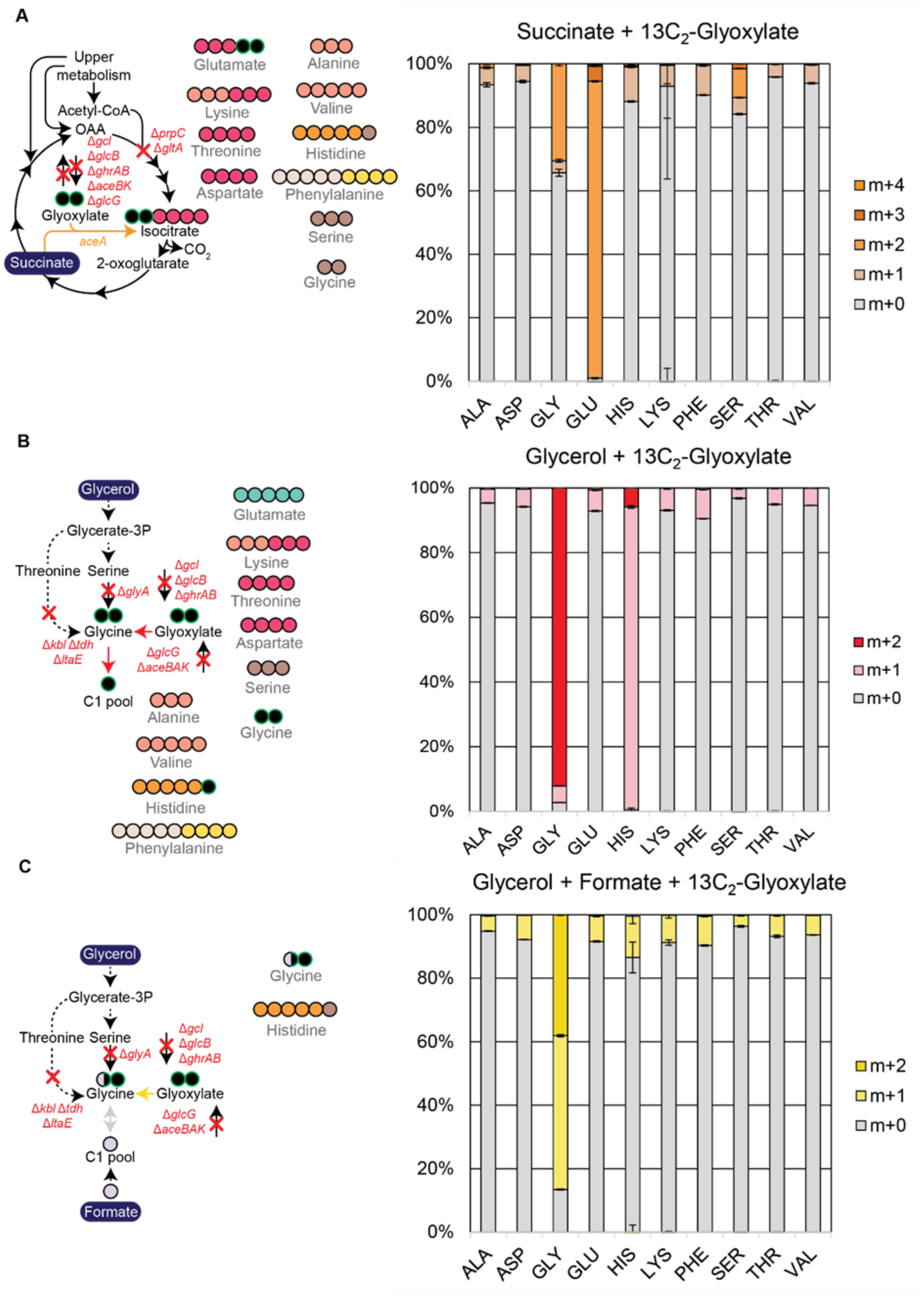


**Fig. S4.**

Labelling patterns for amino acids of metabolic sensor LOW-AUX, UPP-AUX, and TCA-AUX strains grown with $^{13}C_2$ glyoxylate. A) Labelling patterns measured for the 2OXO-AUX strain grown with succinate and $^{13}C_2$ glyoxylate. B) Labelling patterns measured for the C1+GLY-AUX strain grown with glycerol and $^{13}C_2$ glyoxylate. C) Labelling patterns measured for the GLY-AUX strain grown with glycerol and $^{13}C_2$ glyoxylate. The expected patterns for all amino acids except histidine are the same as for the C1+GLY-AUX strain.

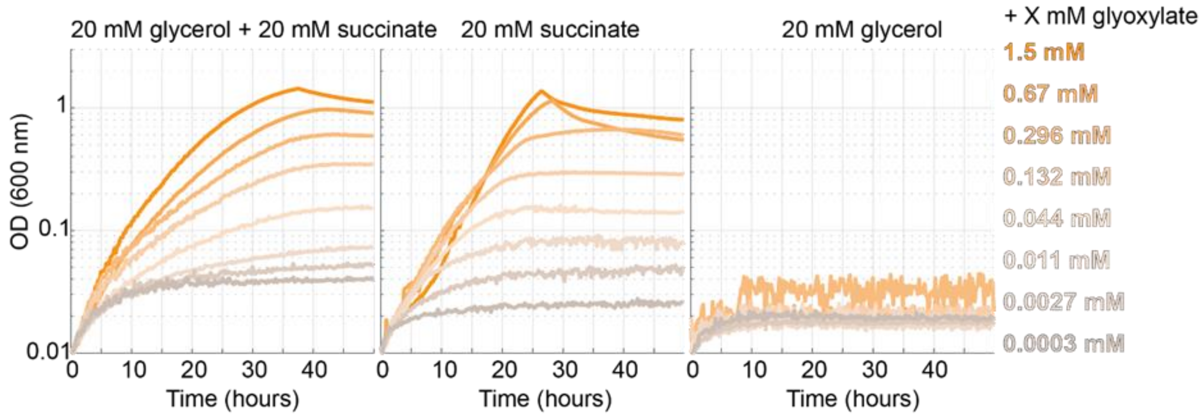


**Fig. S5.**
Additional characterization of the 2OXO-AUX strain. Comparison of glycerol or succinate (+ glyoxylate) growth profiles for the 2OXO-AUX strain to determine the requirement for succinate as co-substrate.

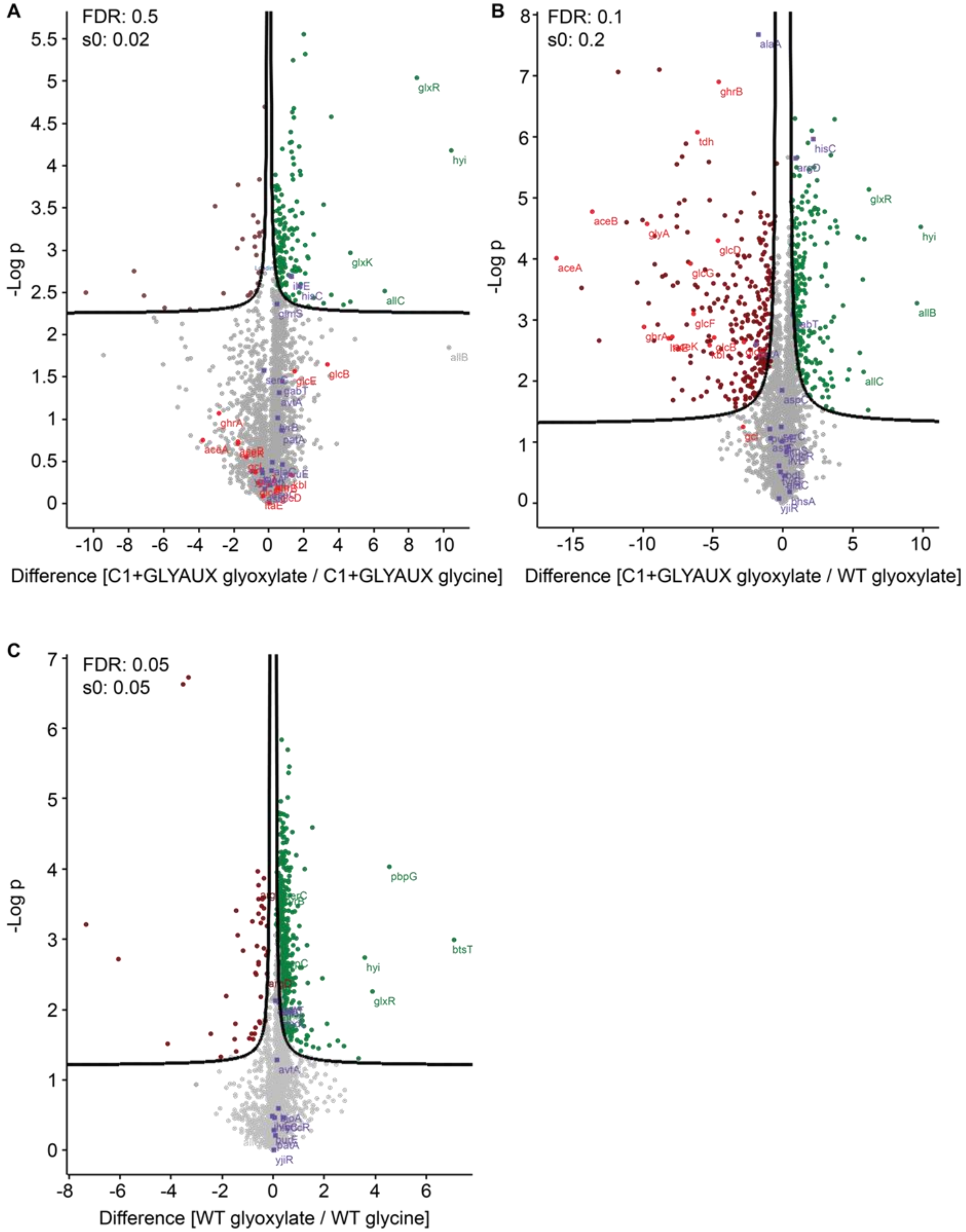


**Fig. S6.**

Proteomics to explore putative transaminase candidates converting glyoxylate to glycine in C1+GLY-AUX. Transaminases are shown in violet with displayed gene names, significantly upregulated genes are shown in green, significantly downregulated ones in brown. Deleted genes are shown in red with displayed gene names. A) Comparison of C1+GLY-AUX grown with 20 mM glycerol + 1 mM glyoxylate vs C1+GLYAUX grown with 20 mM glycerol + 5 mM glycine. HisC and IlvE are significantly upregulated. B) Comparison of C1+GLY-AUX grown with 20 mM glycerol + 1 mM glyoxylate vs a wild type grown with 20 mM glycerol + 1 mM glyoxylate.

HisC and ArgD are significantly upregulated. C) Comparison of wild type grown with 20 mM glycerol + 1 mM glyoxylate vs a wild type grown with 20 mM glycerol + 5 mM glycine.

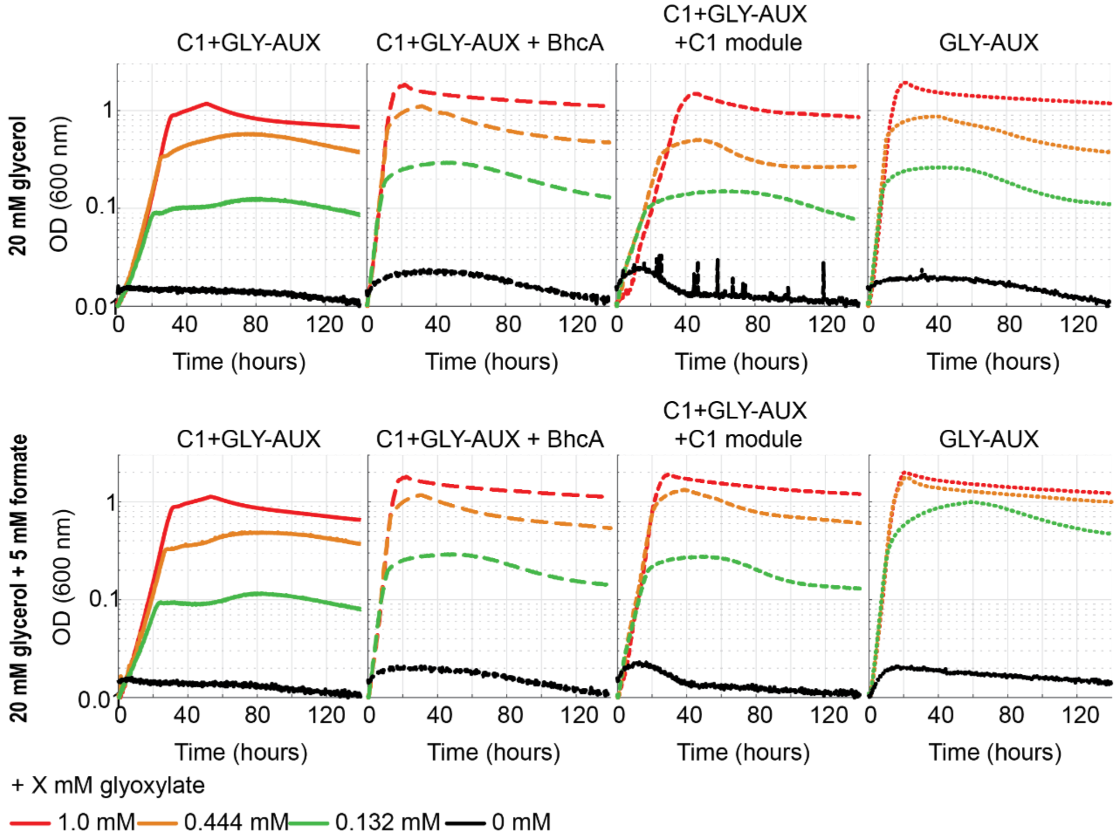


**Fig. S7.**
Additional characterization of C1+GLY-AUX and GLY-AUX strains. The graphs depict the growth profile for the C1+GLY-AUX and GLY-AUX strains under different conditions. From left to right, the panels include: C1+GLY-AUX strain; C1+GLY-AUX strain with the additional heterologously expressed transaminase *bhcA* from *Paracoccus denitrificans*; C1+GLY-AUX with heterologous expression of the C1 module for the reductive glycine pathway (supporting formate to 5,10-methylene-THF conversion); GLY-AUX with the heterologously expressed C1 module and *bhcA*. Top panel: Cultivation under different glyoxylate concentration using glycerol as only co-substrate. Bottom panel: Here, formate is additionally included as co-substrate to support flux via the C1 module of the reductive glycine pathway. *bhcA* and the C1 module are genomically integrated in all strains.

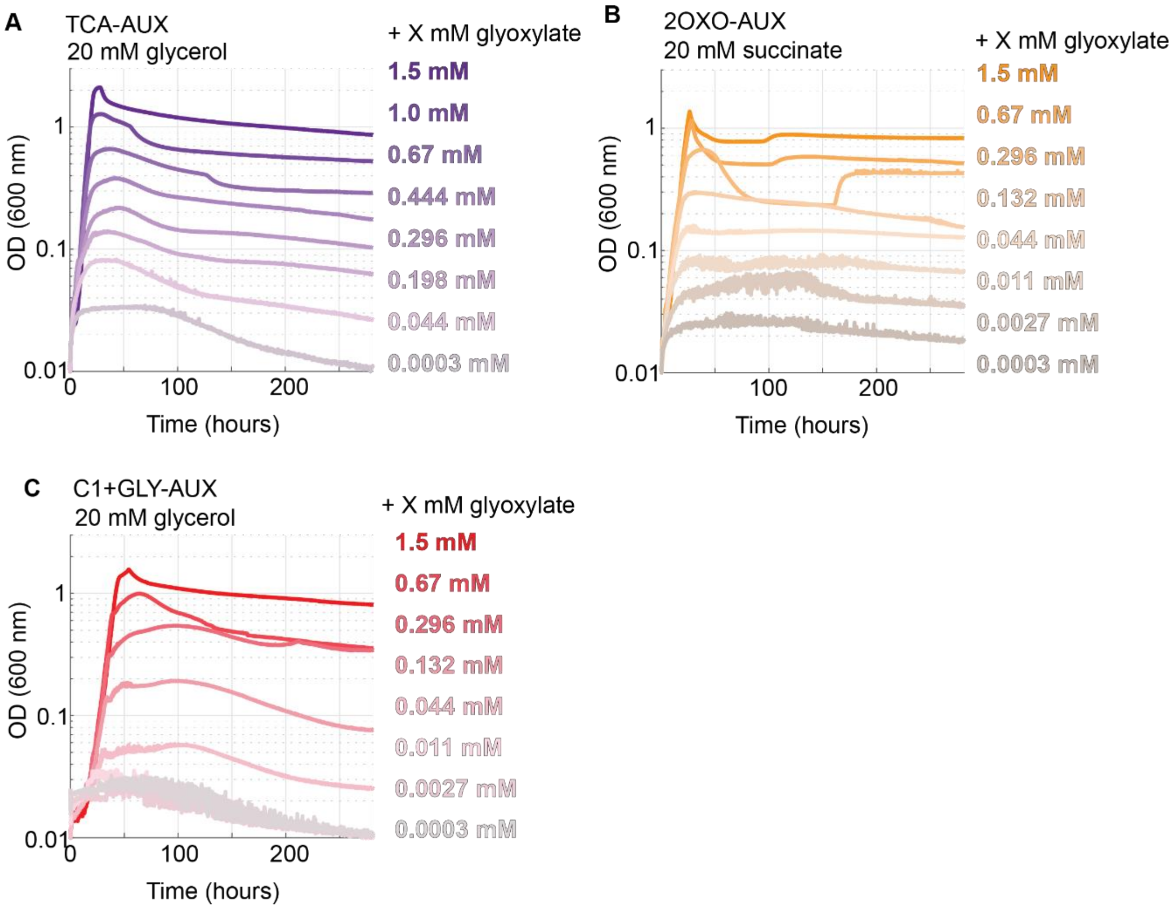


**Fig. S8.**
A-C) Extended growth experiment for more than 250 hours for the three strains to determine the robustness of the auxotrophic phenotype.

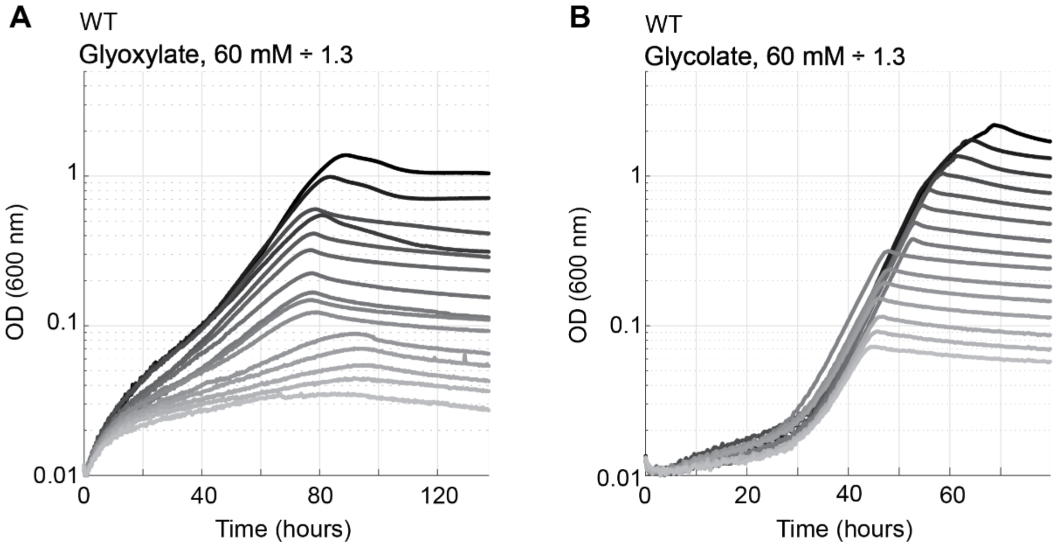


**Fig. S9.**
Growth of a wild type *Escherichia coli* SIJ488 on A) glyoxylate or B) glycolate as sole carbon and energy source. Glycolate concentrations ranged from 60 mM to 0 mM using a 1.3 dilution factor.

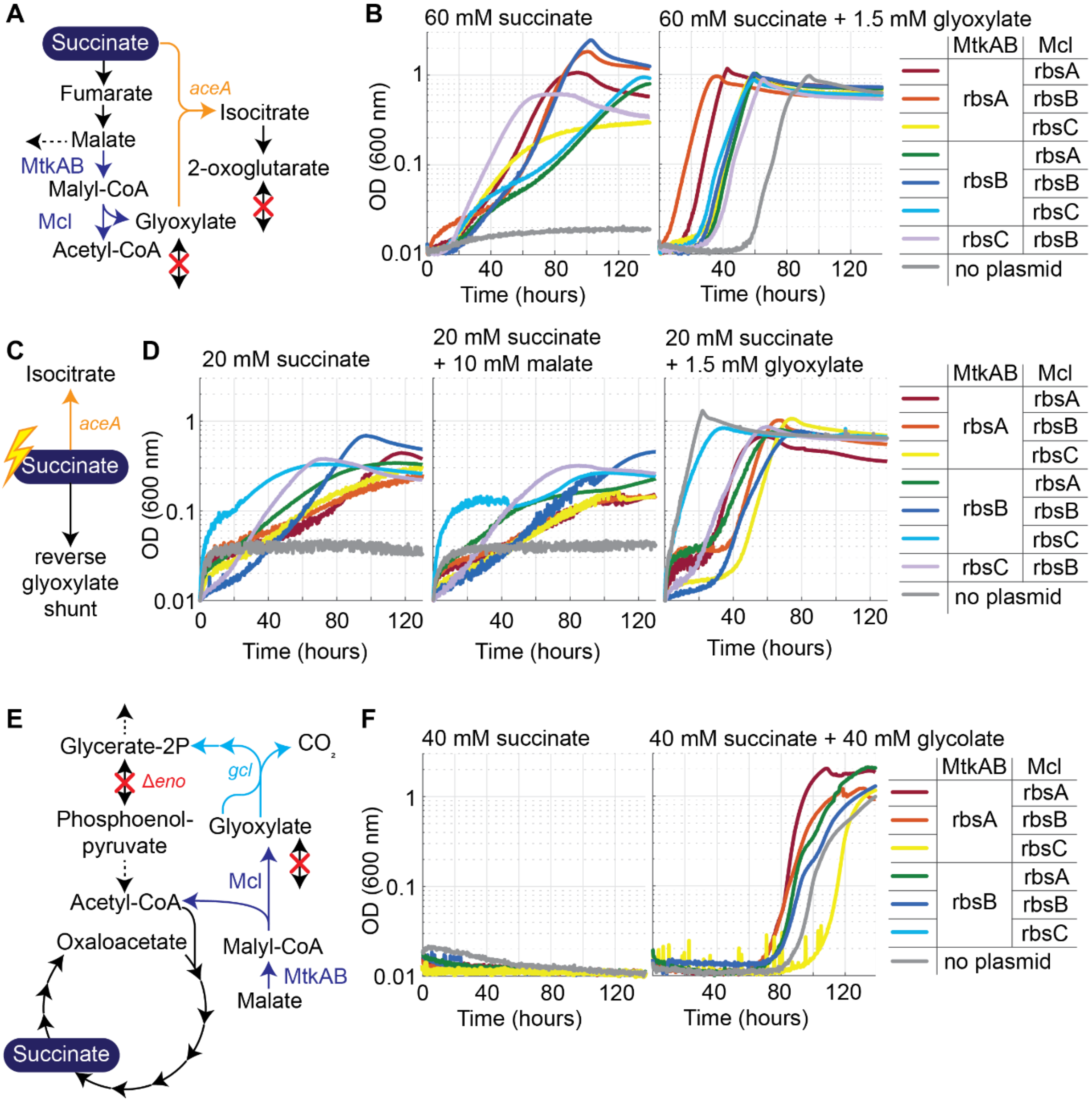


**Fig. S10.**

Use of 2OXO-AUX and UPP-AUX as selection strains for testing MtkAB and Mcl activity under different ribosome binding (RBS) site strengths. A) Selection scheme of 2OXO-AUX strain including MtkAB and Mcl. B) The different behavior of the strains in terms of growth-rates and final biomass concentrations is probably due to a competition for succinate between the MtkAB-Mcl branch and the AceA branch C). D) Even when additional malate was included in the medium, difference in final optical densities -and hence biomass concentrations- could not be prevented. E) Selection scheme for UPP-AUX producing MtkAB and Mcl. F) The behavior of the strain under selective conditions suggests that there is not enough flux capacity for any of the constructs to support growth.

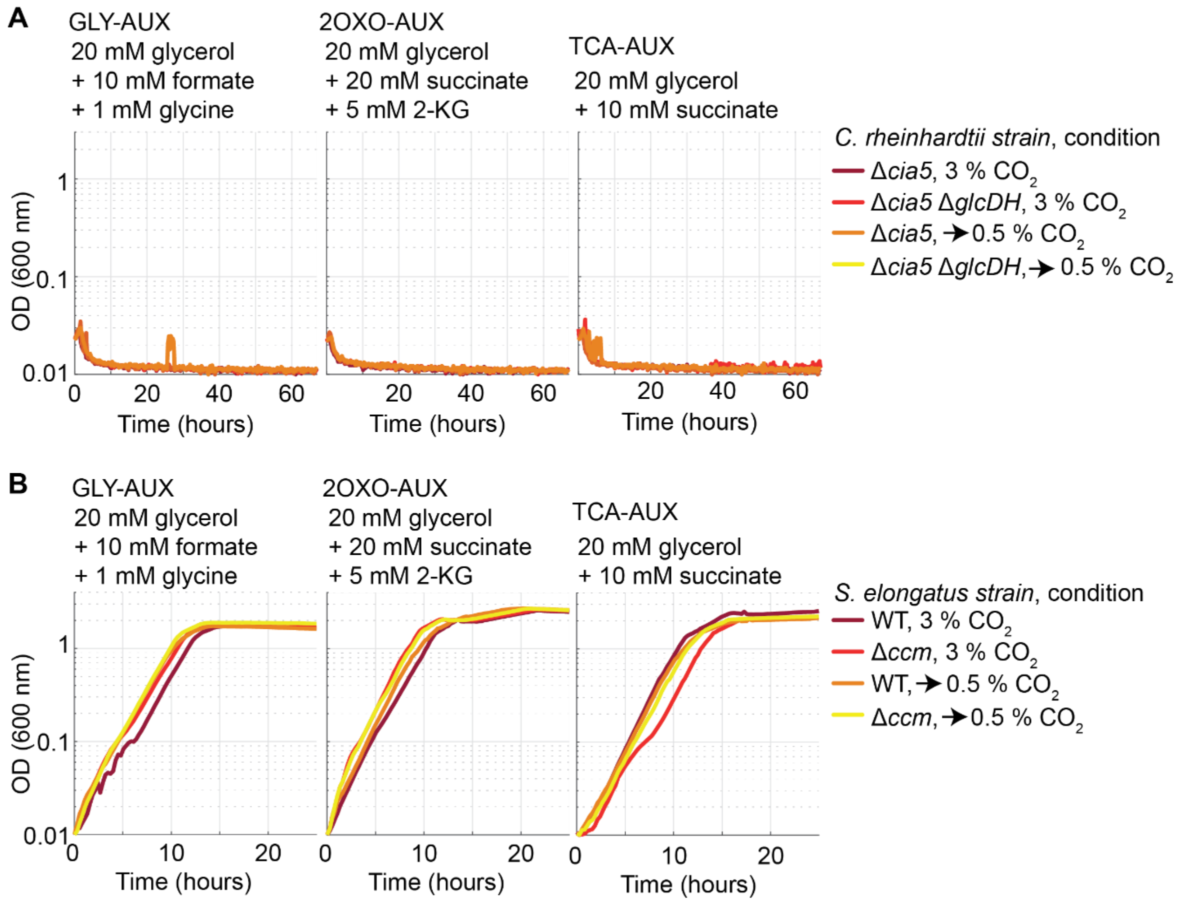


**Fig. S11.**
**Use of the metabolic sensors for determining glycolate in spent fermentative medium.** A, B) Growth profile of GLY-AUX, 2OXO-AUX and TCA-AUX in the cultivation medium of *Chlamydomonas reinhardtii* and *Synechococcus elongatus*, respectively. For *C. rheinhardtii*, we tested knockout strains of the pyrenoid master regulator *cia5* and the pyrenoid master regulator and glycolate dehydrogenase *glcDH.* These strains were grown with acetate in a medium on 3 % $CO_2$, or first on 3 % $CO_2$ and then transferred to 0.5 % $CO_2$ as indicated by the legend to the right. For *Synechococcus elongatus* , we tested a wild type or a pyrenoid (ccm) knockout strain. These strains were grown on 3 % $CO_2$, or first on 3 % $CO_2$ and then moved to 0.5 % $CO_2$ as indicated by the legend to the right.

**Table S1.**

Five chose designs suggested by the in silico search for deletions. The glyoxylate to biomass ratio (GBR) is calculated by dividing maximal growth rate of the strain by the glyoxylate uptake rate. The normalized GBR is calculated by dividing the GBR to the one of a wild-type strain growing on glyoxylate as only carbon source.

| **Strain** | **Knockouts** | **Carbon sources** | **GBR** | **Normalized GBR** |
|---|---|---|---|---|
| 2OXOAUX | ΔCS | Succinate + glyoxylate | 1.1 | 0.02 |
| C1+GLYAUX | ΔICL ΔGHMT2r | Glycerol + glyoxylate | 2.2 | 0.04 |
| TCAAUX | ΔICL ΔPPC | Glycerol + glyoxylate | 3 | 0.05 |
| UPPAUX | ΔICL ΔENO | Succinate + glyoxylate | 8.3 | 0.13 |
| LOWAUX | ΔTPI | Glycerol + glyoxylate | 27.1 | 0.43 |

**Table S2.**

Strains and plasmids used in this study.

| Name | Genetic modifications | Reference |
|---|---|---|
| Wildtype (E. coli SIJ488) | - | (Jensen et al., 2015) |
| NEB5α (E. coli DH5α) | fhuA2Δ(*argF*-lacZ)U169 *phoA glnV*44 Φ80Δ(*lacZ*)M15 *gyrA*96 *recA*1 *relA*1 *endA*1 *thi*-1 *hsdR*17 | #C2987, NEB |
| KEIO Δ*glcB* JW2943 (E. coli MG1655) | Δ*glcB*:KanR | (Baba et al., 2006) |
| 2OXOAUX - glyoxylate only | Δ*gcl* Δ*aceBAK* Δ*glcGB* Δ*gltA* Δ*sucAB* Δ*ghrB* Δ*ghrA* Δ*prpC* SS9:*aceA*:KanR | This work |
| C1+GLYAUX - glyoxylate only | Δ*gcl* Δ*aceBAK* Δ*glcDEFGB* Δ*glyA* Δ*ltaE* Δ*ghrB* Δ*ghrA* Δ*kbl-tdh* | This work |
| TCAAUX - glyoxylate only | Δ*gcl* Δ*aceBAK* Δ*glcDEFGB* Δ*ghrB* Δ*ghrA* Δ*ppc* Δ*maeA* Δ*pck* Δ*maeB* SS9:*glcB* | (Claassens et al., 2020) |
| C1+GLYAUX (+ BhcA) - glyoxylate only | Δ*gcl* Δ*aceBAK* Δ*glcDEFGB* Δ*glyA* Δ*ltaE* Δ*ghrB* Δ*ghrA* Δ*kbl-tdh* Δ*aceBAK*::Prom.S-rbsB-BhcA::CAP (UniProt A1B8Z3) | This work |
| GLYAUX (endogenous transamination) - glyoxylate only | Δ*gcl* Δ*aceBAK* Δ*glcDEFGB* Δ*glyA* Δ*ltaE* Δ*ghrB* Δ*ghrA* Δ*kbl-tdh* SS9::Prom.S-rbsC:MeFtfL-MeFch-MeMtdA (Kan) | This work |
| GLYAUX - glyoxylate only | Δ*gcl* Δ*aceBAK* Δ*glcDEFGB* Δ*glyA* Δ*ltaE* Δ*ghrB* Δ*ghrA* Δ*kbl-tdh* Δ*aceBAK*::Prom.S-rbsB-BhcA::CAP (UniProt A1B8Z3); SS9::Prom.S-rbsC:MeFtfL-MeFch-MeMtdA (Kan) | This work |
| GLYAUX | Δ*gcl* Δ*aceBAK* Δ*glcGB* Δ*glyA* Δ*ltaE* Δ*ghrB* Δ*ghrA* Δ*kbl-tdh* Δ*aceBAK*::Prom.S-rbsB-BhcA::CAP (UniProt A1B8Z3); SS9::Prom.S-rbsC:MeFtfL-MeFch-MeMtdA (Kan), WT *glcDEF* | This work |
| C1+GLYAUX | Δ*gcl* Δ*aceBAK* Δ*glcGB* Δ*glyA* Δ*ltaE* Δ*ghrB* Δ*ghrA* Δ*kbl-tdh* WT *glcDEF* | This work |
| TCAAUX | Δ*gcl* Δ*aceBAK* Δ*glcGB* Δ*ghrB* Δ*ghrA* Δ*ppc* Δ*maeA* Δ*pck* Δ*maeB* WT *glcDEF* | This work |
| UPPAUX | Δeno Δgcl ΔghrAB ΔaceBA | This work |
| LOWAUX | ΔtpiA ΔmgsA | This work |
| **Plasmid** | **Notes** | **Reference** |
| pZ-ASS-rbsA- | Expression plasmid with p15A origin of replication and S-promoter for the expression of Malate thiokinase (expression order rbsA-MtkB-rbsA-MtkA) from *Methylococcus capsulatus* | This work |

| | | |
|---|---|---|
| MtkAB-rbsA-Mcl | and the codon-optimized malyl-CoA lyase (rbsA-Mcl) from *Rhodobacter sphaeroides*. | |
| pZ-ASS-rbsA-MtkAB-rbsB-Mcl | Expression plasmid with p15A origin of replication and S-promoter for the expression of Malate thiokinase (expression order rbsA-MtkB-rbsA-MtkA) from *Methylococcus capsulatus* and the codon-optimized malyl-CoA lyase (rbsA-Mcl) from *Rhodobacter sphaeroides*. | This work |
| pZ-ASS-rbsA-MtkAB-rbsC-Mcl | Expression plasmid with p15A origin of replication and S-promoter for the expression of Malacte thiokinase (expression order rbsA-MtkB-rbsA-MtkA) from Methylococcus capsulatus and the codon-optimized malyl-CoA lyase (rbsA-Mcl) from Rhodobacter sphaeroides. | This work |
| pZ-ASS-rbsB-MtkAB-rbsA-Mcl | Expression plasmid with p15A origin of replication and S-promoter for the expression of Malate thiokinase (expression order rbsA-MtkB-rbsA-MtkA) from *Methylococcus capsulatus* and the codon-optimized malyl-CoA lyase (rbsA-Mcl) from *Rhodobacter sphaeroides*. | This work |
| pZ-ASS-rbsB-MtkAB-rbsB-Mcl | Expression plasmid with p15A origin of replication and S-promoter for the expression of Malate thiokinase (expression order rbsA-MtkB-rbsA-MtkA) from *Methylococcus capsulatus* and the codon-optimized malyl-CoA lyase (rbsA-Mcl) from *Rhodobacter sphaeroides*. | This work |
| pZ-ASS-rbsB-MtkAB-rbsC-Mcl | Expression plasmid with p15A origin of replication and S-promoter for the expression of Malate thiokinase (expression order rbsA-MtkB-rbsA-MtkA) from *Methylococcus capsulatus* and the codon-optimized malyl-CoA lyase (rbsA-Mcl) from *Rhodobacter sphaeroides*. | This work |
| pZ-ASS-rbsC-MtkAB-rbsB-Mcl | Expression plasmid with p15A origin of replication and S-promoter for the expression of Malate thiokinase (expression order rbsA-MtkB-rbsA-MtkA) from *Methylococcus capsulatus* and the codon-optimized malyl-CoA lyase (rbsA-Mcl) from *Rhodobacter sphaeroides*. | This work |
| pTE3275 | Template for the amplification of Malate thiokinase (MtkBA). Expression plasmid with p15A origin of replication and pLacO for the expression of several genes, including Malate thiokinase (MtkBA). | (Luo et al., 2023) |
| pTE3262 | Template for the amplification of Malyl-CoA lyase. Expression plasmid with p15A origin of replication and pLacO for the expression of several genes, including Malyl-CoA lyase (MtkBA). | This work |
| pSEVA64MBEC-aceAB,glcB | Base editing plasmid (pSEVA64 backbone) used for the functional deletion of aceA, aceB and glcB | This study |
| pSEVA64MBEC-ghrAB | Base editing plasmid (pSEVA64 backbone) used for the functional deletion of ghrA and ghrB | This study |

**Table S3.**

List of targeting spacers used for functional inactivation of metabolic genes using a CRISPR/Cas9-mediated cytidine deaminase base editor.

| Name | Sequence (5'→3') | Target gene |
|---|---|---|
| sp1_glcB | CAGCCAGGCGGGGCCGCAGC | *glcB* |
| sp2_glcB | TTCAGGCAGCGCTTGATGAG | *glcB* |
| sp1_aceB | GCCACAACGCAATAAACTTC | *aceB* |
| sp2_aceB | TCCTGCCAGGACTGGGTTTT | *aceB* |
| sp1_aceA | TGGACTCAACCGCGTTGGGA | *aceA* |
| sp2_aceA | CGATCAGATCCAATGGTCCG | *aceA* |
| sp1_ghrB | CTGGCACAACGTGCGCACTT | *ghrB* |
| sp1_ghrA | CACCATTGGGTATCGAACGT | *ghrA* |
| sp2_ghrA | TCCAGCAAAATAGTTCGCAT | *ghrA* |

**Table S4.**

Primers used in this study. Purpose (1 = knockout cassette construction, 2 = deletion verification, 3 = integration verification, 4 = cloning).

| Primer Name | Sequence (5' -> 3') | Purpose |
|---|---|---|
| gcl_KO_fwd | AATTTGAAAGTTGGAAAAATTTTCCAATAAATAGAGGTAGGAATAAAATGATTAACCCTCACTAAAGGGCG | 1 |
| gcl_KO_rvs | GCAGAGAAACGTAACATTATTTATCTCCCTTATTCATAGTGCATGAAGCATAATACGACTCACTATAGGGCTC | 1 |
| gcl_ver_fwd | CCGGGCCTTCATCAAGACTG | 2 |
| gcl_ver_fwd | CGCCGGTAAATTGTGCAGCG | 2 |
| aceBAK_Cap_KO_fwd | TCGTTCACAGTGGGGAAGTTTTCGGATCCATGACGAGGAGCTGCACGATGGTGTAGGCTGGAGCTGCTTC | 1 |
| aceBAK_Cap_KO_rvs | TGCGGAGAAAAATTATATGGAAGCTTTACTCAAAAAAGCATCTCCCCATAGGAATTAGCCATGGTCCATATG | 1 |
| aceBAK_ver_fwd | TCCGAAACGTACCTCAGCAG | 2 |
| aceBAK_ver_rvs | GCTTTATCTGACCATGCACGC | 2 |
| glcDEFGB_Cap_KO_fwd | TAAAACTTGCCGGGGCTTTTTGACGCTATTAATGACTTTCTTTTTCGCGGTGTAGGCTGGAGCTGCTTC | 1 |
| glcDEFGB_Cap_KO_rvs | CGCGCAAAAATCAGCTGCCACACAACACAACAAAGCGAAGCCTACTCATGGGAATTAGCCATGGTCCATATG | 1 |
| glcBGFED_ver_fwd | CGGGTAAGGGTTTCTGTTCCGCAC | 2 |
| glcBGFED_ver_rvs | CACACAGTCGACGTTCCGAGGGAAG | 2 |
| ghrB_KO_fwd | CATATTTCAGGCTAAGGTGATCGCCTTATCAGTGAATGGAGAGAAGCATGATTAACCCTCACTAAAGGGCG | 1 |
| ghrB_KO_rvs | ATCGGGCTTTACTCTACGCAGTCGCGGCTTAGTCCGCGACGTGCGGATTCTAATACGACTCACTATAGGGCTC | 1 |
| ghrB_ver_fwd | GGGCCTTGGCCCGGCTAACC | 2 |
| ghrB_ver_rvs | GCTTTGTCGCAAATGGCATC | 2 |
| ghrA_KO_fwd | AACGATAAGTGCGAATAAATTTCGCACAACGCTTTTCGGGAGTCAGTATGATTAACCCTCACTAAAGGGCG | 1 |
| ghrA_KO_rvs | CCAAGGATAGCAGGAATCCTGATGCTTTATTAGTAGCCGCGTGCGCGGTCTAATACGACTCACTATAGGGCTC | 1 |
| ghrA_ver_fwd | CTGGAACGGGCGCTAATTTAG | 2 |
| ghrA_ver_rvs | GGGCCATGATCGGTGATCGC | 2 |
| gltA_KO_fwd | TAAAATATTTACAACTTAGCAATCAACCATTAACGCTTGATATCGCTTTTAATTAACCCTCACTAAAGGGCG | 1 |
| gltA_KO_rvs | TTTAAGTTCCGGCAGTCTTACGCAATAAGGCGCTAAGGAGACCTTAAATGTAATACGACTCACTATAGGGCTC | 1 |
| gltA_ver_fwd | TGGGTTGCTGATAATTTGAGCTGT | 2 |
| gltA_ver_rvs | GCTCTGTACCCAGGTTTTCCCC | 2 |
| sucA_KO_fwd | AGTATCCACGGCGAAGTAAGCATAAAAAAGATGCTTAAGGGATCACGATGAATTAACCCTCACTAAAGGGCG | 1 |
| sucB_KO_rvs | GGTCTACAGTGCAGGTGAAACTTAAACTACTACACGTCCAGCAGCAGACGTAATACGACTCACTATAGGGCTC | 1 |
| sucA_ver_fwd | CCGGCCTACAAGTCATTACCCG | 2 |
| sucB_ver_rvs | ACCGTAGACCGAATATTTCGTATCGC | 2 |
| prpC_KO_fwd | CCGACAACAATCTCGACCCTACAAATGATAACAATGACGAGGACAACATGAATTAACCCTCACTAAAGGGCG | 1 |
| prpC_KO_rvs | TACGTTTCCTTATTGTTATTCGTAGAGGTTTACTGGCGCTTATCCAGCGCTAATACGACTCACTATAGGGCTC | 1 |

| | | |
|---|---|---|
| prpC_Ver_fwd | ATCCGACAACCGATGCCTGATG | 2 |
| prpC_Ver_rvs | AGTAATGTGCGGTGTCGTAGGC | 2 |
| Ppc_KO_fwd | GAAGGATACAGGGCTATCAAACGATAAGATGGGGTGTCTGGGGTAAT ATGAATTAACCCTCACTAAAGGGCG | 1 |
| Ppc_KO_rvs | AAAGCACGAGGGTTTGCAGAAGAGGAAGATTAGCCGGTATTACGCAT ACCTAATACGACTCACTATAGGGCTC | 1 |
| ppc_ver_fwd | ACGAGGGTGTTAGAACAGAAGT | 2 |
| ppc_ver_rvs | CAAAGCCCGAGCATATTCGC | 2 |
| maeA_KO_fwd | CCCGGTAGCCTTCACTACCGGGCGCAGGCTTAGATGGAGGTACGGCG GTAAATTAACCCTCACTAAAGGGCG | 1 |
| maeA_KO_rvs | GGCCGACGCCCTGGCGGTAAAGCAAAGACGATAAAAGCCCCCCAGG GATGTAATACGACTCACTATAGGGCTC | 1 |
| maeA_ver_fwd | CCTTCGTAACAACTGCAGCC | 2 |
| maeA_ver_rvs | TTAAACACCATTGCCTGCGC | 2 |
| pck_KO_fwd | CTGGATAGATATTCTCCAGCTTCAAATCATTACAGTTTCGGACCAGCC GCAATTAACCCTCACTAAAGGGCG | 1 |
| pck_KO_rvs | AAAGACTTTACTATTCAGGCAATACATATTGGCTAAGGAGCAGTGAA ATGTAATACGACTCACTATAGGGCTC | 1 |
| pck_ver_fwd | ATCTATGAGCCTTGTCGCGG | 2 |
| pck_ver_rvs | GCAGGGCACGACAAAAGAAG | 2 |
| maeB_KO_fwd | TTCAGGGTAAGCGTGAGAGTTAAAAAAAATTACAGCGGTTGGGTTTG CGCAATTAACCCTCACTAAAGGGCG | 1 |
| maeB_KO_rvs | TTGCCCACACACTTTATTTGTGAACGTTACGTGAAAGGAACAACCAA ATGTAATACGACTCACTATAGGGCTC | 1 |
| maeB_ver_fwd | AGAGATATTCGCTGTGGTGCA | 2 |
| maeB_ver_rvs | GCAGACAGGCATGGTATTGC | 2 |
| glyA_KO_fwd | AGCACATTGACAGCAAATCACCGTTTCGCTTATGCGTAAACCGGGTA ACGAATTAACCCTCACTAAAGGGCG | 1 |
| glyA_KO_rvs | CCTATAAAGGCCAAAAATTTTATTGTTAGCTGAGTCAGGAGATGCGG ATGTAATACGACTCACTATAGGGCTC | 1 |
| glyA_ver_fwd | CGCGATAACGTAGAAAGGCT | 2 |
| glyA_ver_rvs | GTAGAAATGGGCGGTTAACT | 2 |
| ltaE_KO_fwd | CTAAAATGCGTTGCGGCACGTCTCTCTCCTTAACGCGCCAGGAATGCA CGAATTAACCCTCACTAAAGGGCG | 1 |
| ltaE_KO_rvs | GGCTAAGGTAAAAAGGGTGGCATTTCCCGTCATAATAAGGACATGCC ATGTAATACGACTCACTATAGGGCTC | 1 |
| ltaE_ver_fwd | TAGCCATTCCACCGCAGATC | 2 |
| ltaE_ver_rvs | AGGATCTGATGCCCTTGCTG | 2 |
| kbl-tdh_KO_fwd | AAGGGCTGGAATACCAGCCCTTGTTCGTGTTAATCCCAGCTCAGAAT AACAATTAACCCTCACTAAAGGGCG | 1 |
| kbl-tdh_KO_rvs | AAGTTTGGGTAATATGTGCTGGAATTTGCCCTGTCTGGAGAATCGCAA TGTAATACGACTCACTATAGGGCTC | 1 |
| kbl-tdh_ver_fwd | CCTCTGAAGTTATCCTGACGTTTTACAGG | 2 |
| kbl-tdh_ver_rvs | CCAACTGAGATAGGAAAAATGATGCTTTGGC | 2 |
| pSS9_total_fwd | GCTTGGTTGAGAATACGCCG | 3 |
| pSS9_total_rvs | GCCTACGATTACGCATGGCTT | 3 |
| 971_ASS_A-MtkB_rvs | tacgctcatTCCAAACCTCCTTAAAGTTAAACAAAATTATTTCTATTAACTA GTGAATTCGGTCAGTG | 4 |
| 973_ASS-B-MtkB_rvs | tacgctcatCTCAGTACCTCCTCATTTTGTTTAAAGTTAAACAAAATTATTT CTATTAACTAGTGAATTCGGTCAGTG | 4 |

| | | |
|---|---|---|
| 974_ASS-C-MtkB_rvs | tacgctcattcttgcctcttaacttTAAAGTTAAACAAAATTATTTCTATTAACTAGTGAATTCGGTCAGTG | 4 |
| 972_ASS-Mcl_fwd | ccgagatgattagtgcttaaGCTAGCGCGGCCGCTCGAGt | 4 |
| Ec864_MtkB-rbsA_fwd | aataattttgtttaactttaAGGAGGTTTGGAatgagcgtattcgttaacaagcac | 4 |
| Ec865_MtkB-rbsB_fwd | aataattttgtttaactttaAACAAAATGAGGAGGTACTGAGatgagcgtattcgttaacaagcac | 4 |
| Ec858_MtkB-rbsC_fwd | aataattttgtttaactttaaagttaagaggcaagaatgagcgtattcgttaacaagcac | 4 |
| Ec859_MtkB-rvs | GGGTTCAGTTACTTAACAGAACCTCCGATCAACGCTAGCTCTAGAtcagaatctgattccgtgtt | 4 |
| Ec866_MtkA-rbsA_fwd | TCTGTTAAGTAACTGAACCCTAATAGAAATAATTTTGTTTAACTTTAAGGAGGTTTGGAATGaatatccatgagtacca | 4 |
| Ec867_MtkA-rbsB_fwd | TCTGTTAAGTAACTGAACCCTAATAGAAATAATTTTGTTTAACTTTAAACAAAATGAGGAGGTACTGAGATGaatatccatgagtacca | 4 |
| Ec860_MtkA-rbsC_fwd | TCTGTTAAGTAACTGAACCCTAATAGAAATAATTTTGTTTAACTTTAAAGTTAAGAGGCAAGAATGaatatccatgagtacca | 4 |
| Ec861_MtkA_rvs | AAGAGGTAAGCGTCACTAACGACAttatcccttgacgatggcga | 4 |
| Ec868_Mcl-rbsA_fwd | GTTAGTGACGCTTACCTCTTTAATAGAAATAATTTTGTTTAACTTTAAGGAGGTTTGGAatgagttttcggttgcaacct | 4 |
| Ec869_Mcl-rbsB_fwd | GTTAGTGACGCTTACCTCTTTAATAGAAATAATTTTGTTTAACTTTAAACAAAATGAGGAGGTACTGAGatgagttttcggttgcaacct | 4 |
| Ec862_Mcl-rbsC_fwd | GTTAGTGACGCTTACCTCTTTAATAGAAATAATTTTGTTTAACTTTAAAGTTAAGAGGCAAGAatgagttttcggttgcaacct | 4 |
| Ec863_Mcl_rvs | ctcgagcggccgcgctagcttaagcactaatcatctcgg | 4 |
| oEO_088 | GGGCCTTGGCCCGGCTAACC | 2 |
| oEO_089 | GCTTTGTCGCAAATGGCATC | 2 |
| oEO_182 | atccatgacgaggagctgc | 2 |
| oEO_183 | aacgaccgcagttcagacc | 2 |
| oEO_184 | caccgaagaaggtatgcgc | 2 |
| oEO_185 | aagtgaactgccgctgcac | 2 |
| oEO_186 | ttgatgtgcgccggatcg | 2 |
| oEO_187 | gcgtcgtgaaactaagggc | 2 |
| oEO_188 | acacactttccaggcgtgctcc | 2 |
| oEO_189 | gcgagcaggtcatcttccac | 2 |
| oEO_190 | agaaaccgtcgccgatacgc | 2 |
| oEO_191 | cagctgatgcattgcaaccg | 2 |

**Table S5.**

Sequences of heterologous genes used in this study. To be included as Excel table.

**Table S6.**
Parameters used for LC-MS/MS based glycolate quantification.

| Name | Precursor Ion | Product Ion | Collision energy [V] | Fragmentor Voltage [V] | Cell Accelerator Voltage [V] | Dwell time [msec] | Polarity |
|---|---|---|---|---|---|---|---|
| Glycolate | 75 | 75 | 0 | 380 | 5 | 150 | Negative |
| Glycolate | 75 | 47 | 9 | 380 | 5 | 150 | Negative |
